# Pan-genome of pear provides insights into the fruit quality traits differentiation between Asian and European pears

**DOI:** 10.1101/2023.09.29.560244

**Authors:** Baopeng Ding, Haifei Hu, Tingting Liu, Muhammad Tahir ul Qamar, Yujing Lin, Ruirui Xu, Zhiwen Chen, Yuqin Song, Guangqi He, Youzhi Han, Huangping Guo, Jun Qiao, Jianguo Zhao, Xinxin Feng, Sheng Yang, Shaofang He, Liulin Li, Rajeev K. Varshney, Xuhu Guo

**Affiliations:** Engineering Research Center of Coal-Based Ecological Carbon Sequestration Technology of the Ministry of Education and Key Laboratory of National Forest and Grass Administration for the Application of Graphene in Forestry, Shanxi Datong University, Datong, Shanxi, 037009, P.R. China; College of Horticulture, Shanxi Agricultural University, Taigu, Shanxi, 030801, P.R. China; Rice Research Institute, Guangdong Academy of Agricultural Sciences & Key Laboratory of Genetics and Breeding of High Quality Rice in Southern China (Co-construction by Ministry and Province), Ministry of Agriculture and Rural Affairs & Guangdong Key Laboratory of New Technology in Rice Breeding & Guangdong Rice Engineering Laboratory; Integrative Omics and Molecular Modeling Laboratory, Department of Bioinformatics and Biotechnology, Government College University Faisalabad (GCUF), Faisalabad, 38000, Pakistan; College of Biology and Oceanography, Weifang University, Weifang, Shandong, 261061, P.R. China; State Key Laboratory of Plant Physiology and Biochemistry, College of Life Sciences, Zhejiang University, Hangzhou, Zhejiang, 310058, P. R. China; College of Forestry, Shanxi Agricultural University, Taigu, Shanxi, 030801, P.R. China; Pomology Institute, Shanxi Agricultural University, Taigu, Shanxi, 030801, P.R. China; Wuhan Tanma Technology and Shanxi Editor Technology, Wuhan&Taiyuan, Hubei &Shanxi, 430206 & 030000, P.R. China; Centre for Crop & Food Innovation, State Agricultural Biotechnology Centre, Food Futures Institute, Murdoch University, Western Australia, Perth, 6000, Australia; School of Life Sciences, Shanxi Datong University, Datong, Shanxi, 037009, P.R. China

**Keywords:** Phased diploid genome, Pangenome, PAV, pear, fruit quality traits

## Abstract

The pear (*Pyrus spp*.) is a remarkable fruit, well known for its diverse flavors, textures, culinary versatility, and global horticultural importance. However, the genetic diversity responsible for its extensive phenotypic variations remains largely unexplored. Here, we *de novo* assembled and annotated the genomes of the maternal (PsbM) and paternal (PsbF) lines of the hybrid ‘Yuluxiang’ pear and constructed the first pear pangenome of 1.15Gb by combining these two genomes with five previously published pear genomes. Using the constructed pangenome, we identified 21,224 gene PAVs and 1,158,812 SNPs in the non-reference genome that were absent in the PsbM reference genome. Compared with SNP markers, we found that PAV-based analysis provides additional insights into the pear population structure. In addition, we also revealed that some genes associated with pear fruit quality traits have differential occurrence frequencies and differential gene expression between Asian and European populations. Moreover, our analysis of the pear pangenome revealed a mutated SNP and an insertion in the promoter region of the gene *PsbMGH3.1* potentially enhances sepal shedding in ‘Xuehuali’ which is vital for pear quality. This research helps further capture the genetic diversity of pear populations and provides valuable genomic resources for accelerating pear breeding.

## Introduction

The pear (*Pyrus spp.*), an illustrious member of the Rosaceae family, holds global renown for its palatable taste, nutritional offerings, and significant economic implications (Wang et al., 2023a). Pear is originally from China and has diverse germplasm resources and a long cultivation history. Currently, at least 22 distinct pear species are recognized, with over 5,000 accessions preserved by researchers globally (Wu et al., 2018). This collection spans a myriad of morphologies, a spectrum of physiological differences, and broad adaptability to diverse ecological niches. Within this array of varieties, the ‘Yuluxiang’ pear emerges as a breeding marvel, representing three dedicated breeder generations from the Fruit Research Institute at Shanxi Agricultural University’s Pear Research Group (Ding et al., 2021; Wu et al., 2019). Accredited by the Ministry of Agriculture, this pear ascends in popularity, characterized as a mid-maturing, shelf-stable red pear. Its distinguishing traits include a robust size, inviting fruity aroma, vibrant red hue, minimal core, rounded contour, and a succulent, crisp bite. Given these attributes, the ‘Yuluxiang’ pear merits its status as a pivotal genetic material, elucidating its morphology, biology, and genetic nuances.

As bioinformatics assembly algorithms advanced and genome sequencing costs decreased, *Pyrus* genomics studies have proliferated, yielding five distinct genome assemblies from varied pear types. This includes the European cultivar pear (*P. communis*) (Chagne et al., 2014), the Chinese white pear (*P. bretschneideri*) (Wu et al., 2013), a Chinese wild pear (*P. betuleafolia*) (Dong et al., 2020), a Japanese pear (*P. pyrifolia*) (Shirasawa et al., 2021), and a dwarfing pear [(Pyrus ussuriensis × communis) × spp.] (Ou et al., 2019). Among these, two genomes have undergone successful assembly and haplotype separation. Yet, the currently published pear genomes are required to be further improved, especially in aspects of genome completeness and contiguity. High-quality pear genomes facilitate further studies into the pear’s evolutionary history, adaptability, and potential for genetic improvement. Long-read sequencing technology such as PacBio HiFi (Cheng et al., 2021) is required to produce a high-quality genome by enabling the assembly of complex regions, repetitive sequences, and structural variations, leading to a more accurate and comprehensive representation of the genome’s architecture and functional elements.

Although several genomes from the *Pyrus* genus have been published, the genetic diversity within the genus cannot be sufficiently represented by a single species genome, potentially leaving crucial agronomic traits in non-reference genome regions unexplored. Consequently, constructing a pan-genome is imperative, enabling the encapsulation of the full spectrum of genetic variations within a species. The dramatic reduction in sequencing costs has facilitated the application of high-throughput sequencing technologies to pan-genomes, as evidenced by the first soybean pan-genome (Li et al., 2014). Recently, the focus on pangenomics has surged, encompassing a range of plants, from crops to ornamentals and trees. This approach has expanded from staple crops such as wheat (Walkowiak et al., 2020), rice (Qin et al., 2021; Wang et al., 2023b), and potatoes (Tang et al., 2022) to oil crops such as soybeans (Bayer et al., 2022; Liu et al., 2020), rapeseed (Song et al., 2020), and sesame (Yu et al., 2019). It also covers legumes such as peas (Yang et al., 2022), pigeon pea (Zhao et al., 2020) and chickpeas (Varshney et al., 2021), horticultural plants such as apple (Sun et al., 2020), tomatoes (Zhou et al., 2022), and extends to tree species such as *Amborella* (Hu et al., 2022), pecan (Lovell et al., 2021) and yellowhorn (Wang et al., 2023c). The growing emphasis on pan-genomic research is evident and poised to continue (Tahir et al., 2020).

In this context, our study assembled two high-quality pear genomes from the maternal (PsbM) and paternal (PsbF) lines of the hybrid ‘Yuluxiang’ pear, integrating PacBio and Hi-C sequencing datasets. Furthermore, we constructed the first pear pangenome by merging the PsbM genome with the PsbF genome and other previously released pear genomes, offering a more comprehensive view of pear genetic resources. Leveraging this newly constructed pangenome, we revisited the short-read sequencing data from 113 varied pear species, and pinpointed a plethora of gene PAVs which might be linked to pear quality traits, revealing differential prevalence between Asian and European pear groups.

## Results

### Chromosome-length assemblies of pear genomes

Using the PacBio CCS (HiFi) reads and Illumina short reads, we successfully *de novo* assembled two parent genomes of the hybrid ‘Yuluxiang’ pear cultivar from Taigu of Jinzhong city, China’s Shanxi, including the paternal line PsbF (*P. sinkiangensis*) from Korla city, China’s Xinjiang, and the maternal line PsbM (*P. bretschneideri*) from Shijiazhuang city, China’s Hebei (**Figure 1A**). Following correction for misjoins, the contigs were ordered, oriented, and anchored to chromosomes using Hi-C sequencing (**Figure 1 B, C and D**). This allowed the anchoring of approximately 96.2% and 98.8% of sequences to chromosomes for PsbF and PsbM, respectively. Both genome assemblies demonstrated near gapless qualities, with ten of seventeen chromosomes comprising only one contig and the remaining chromosomes containing fewer than three contigs (**Table S1)**. Centromere and telomere-specific repeats were identified across almost all chromosomes in both the PsbF and PsbM genomes (**Table S2**), indicative of the high-quality assembly of these genomes. Moreover, at least 44.5% of repeat sequences were identified within the assembled PsbF and PsbM genomes, with long terminal repeat (LTR) retrotransposon elements being the most prevalent class of transposable elements (TEs), accounted for at least half of the TEs (**Table S3**). The assembly lengths of the PsbF and PsbM genomes are around 538 Mb and 510 Mb, respectively. Both genome assemblies have a contig N50 over 26.6 Mb, which is higher than previously published pear genomes (**Figure 1E**). Benchmarking Universal Single-Copy Orthologs (BUSCO) and Cluster of Essential Genes (CEG) evaluation confirmed the completeness of the PsbF and PsbM genomes to exceed 96% and 98% (**Table S4)**. These results illustrate a high level of contiguity and completeness of our assembled pear genomes.

**Figure 1.**
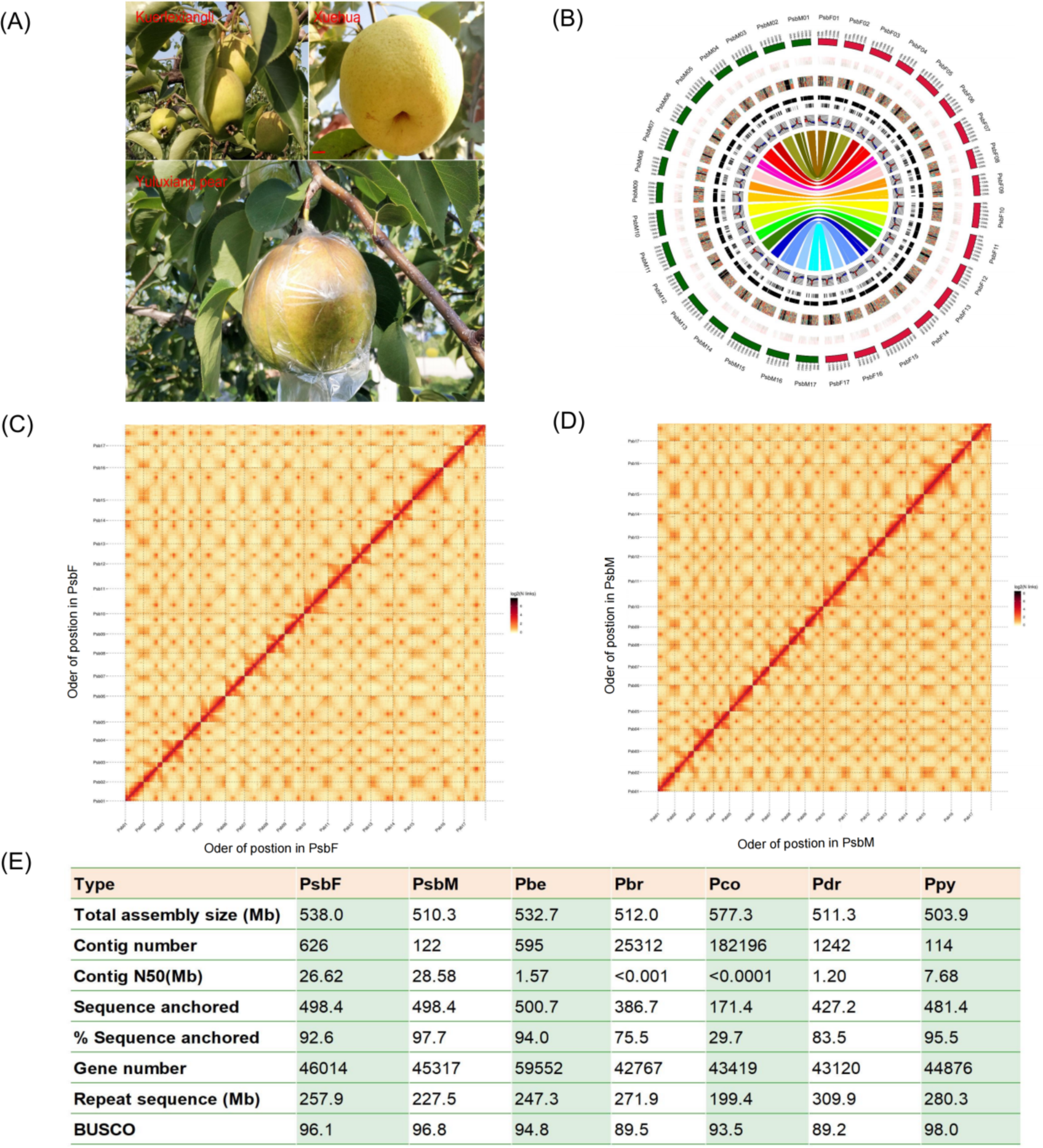
Genome assembly of PsbM and PsbF pear genomes. ‘Yuluxiang’ Pear and Its maternal Kuerlexiangli (PsbF) and paternal Xuehuali(PsbM). (B) Genome-wide characteristics of the PsbM and PsbF pear genomes. (C) and (D) Chromosome contact maps across the seven chromosome-length scaffolds of the PsbM and PsbF pear genomes. (E) Genome assembly and annotation statistics for the PsbM and PsbF pear genome assemblies compared to a Chinese wild pear (Pbe-SD, Pbe), a Japanese pear (Ppy), the Chinese white pear (DSHS, Pbr), the European cultivar pear (Bartlett,Pco) and a dwarfing pear (Zhonggai1, Pdr).

Through a combination of RNA-seq transcript mapping, *ab initio* prediction, and homologous protein searches, we annotated 46,014 and 45,317 gene models in the PsbF and PsbM assemblies, respectively (**Figure 1E**), wherein around 90% of genes garnered functional annotation from at least one functional protein database (**Table S5**). When examined from a genome-wide perspective, gene models exhibited greater density in chromosome arms with fewer repeat elements. Additionally, we annotated 8,514 and 7,402 non-coding RNAs, including rRNAs, tRNAs, miRNAs, and snRNAs, in the PsbF and PsbM genomes, respectively (**Table S6**).

### Large genomic variations in seven pear genomes

The high-quality pear enables the discovery of structural variations (SVs) that are equal to or larger than 50 bp in size. Based on the PsbM genome, we identified SVs in six other pear genomes including a European wild pear (*P. communis,* Pco) (Chagne et al., 2014), a Chinese white pear (*P. bretschneideri*) (Wu et al., 2013), a Chinese wild pear (*P. betuleafolia*, Pbe) (Dong et al., 2020), a Japanese pear (*P. pyrifolia*, Ppy) (Shirasawa et al., 2021), and a dwarfing pear [(*Pyrus ussuriensis* × *communis*) × spp., Pdr] (Ou et al., 2019). In total, 44,681 non-redundant SVs, including 247 inversions, 814 translocations, 431 duplications, 14,837 deletions, and 27,979 insertions were identified, with translocations and inversions having longer sequence lengths than deletions and insertions (**Figure 2A**). Interestingly, compared with other pear genomes, Pdr and Ppy contained the largest number of SVs (**Table S7**). Furthermore, we detected 6,341 high-confident SVs across at least two genomes, encompassing 7 inversions, 15 translocations, 3,989 deletions, and 2,330 insertions (**Table S8**). We also found 319 high-confident SVs located in the gene exon regions that will potentially be affecting the gene functions (**Table S9**).

**Figure 2.**
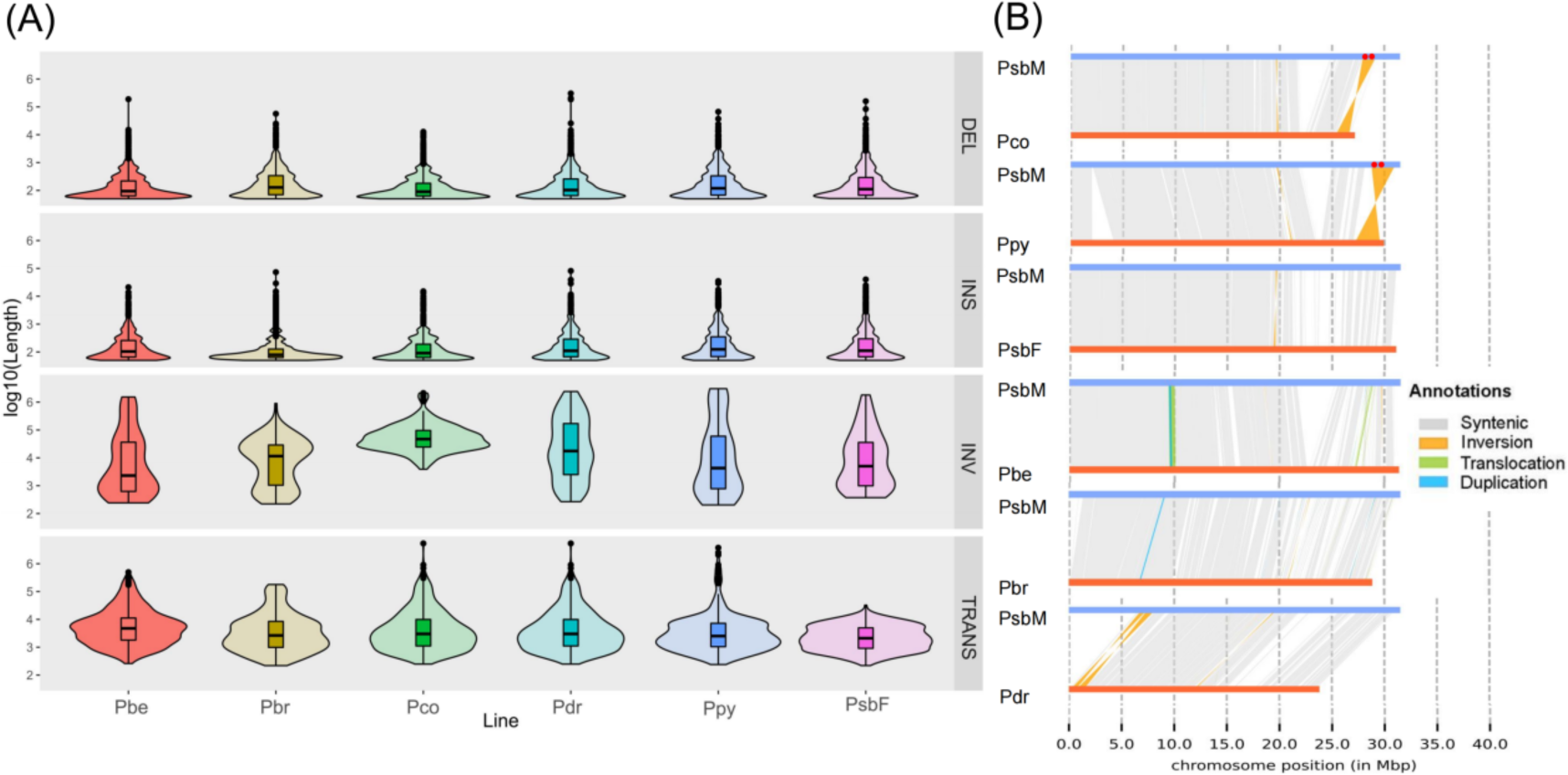
Structural variations across seven pear genome assemblies. (A) The size of SVs (insertions, deletions, translocations and inversions) by comparing six different genomes to the PsbM genome. (B) Synteny map showing the structure of Chromosome 13. Red dots between 25 Mbp to 30 Mbp show the position of two proline-serine-threonine phosphatase-interacting protein genes (PsbM013G02206 and PsbM013G02206) functionally associated with cell division.

Previous studies suggest that inversions are associated with local recombination suppression, facilitating the selection of adaptive traits (Crow et al., 2020; Hamala et al., 2021). For instance, an inversion spanned 646 kb, located in chromosome 13 (PsbM13:19,118,031-19,764,319), was present in Ppy and Pdr genomes but absent in PsbM, PsbF, Pco, Pbr, and Pbe genomes (**Figure 2B**). This inverted region contains 27 gene models in the PsbM genome, including two proline-serine-threonine phosphatase-interacting protein genes (*PsbM013G02206* and *PsbM013G02206*) functionally associated with cell division and chromosome partitioning and located in the inversion breakpoint at around 119Kb distance from the distal breakpoint (**Table S9**). Moreover, deletions and insertions were reported to be related to adaptation to varying pathogen pressures within different growing environments (Dolatabadian et al., 2022; Hu et al., 2022; Zmienko et al., 2014). In this study, we identified that 2 insertions and 9 deletions were overlapped with the exon regions of genes which were functionally associated with disease resistance (**Table S9**). For example, a 663 bp deletion leading to three exons absent was identified in the gene *PsbM006G00203* (homology of *Arabidopsis Disease resistance protein RPM1*) and a 9,393 bp deletion resulting in complete loss of two *Arabidopsis* disease resistance protein homologous genes (*PsbM015G01503* and *PsbM015G01504*) among the Pco, Pdr, and PsbF genomes.

### Construction of the pear pangenome

Using the pangenome construction method similar to the pangenomics studies of *B.napus* (Song et al., 2020), *B.oleracea* (Golicz et al., 2016), and soybean (Torkamaneh et al., 2021), we constructed the first linear pear pangenome by adding the PAV sequences from other six pear genomes to the PsbM reference genome. The constructed pear pangenome is 1.15Gb in size, containing an additional 635 Mb non-reference sequences and hosting 21,560 newly annotated high confidence genes which were missing in the reference genome. A higher percentage (76.0%) of repetitive elements accounted for the non-reference genome than the reference genome (54.3%) (**Table S10**), indicating that transposable elements might be the important driver of presence/absence variations. By analyzing the read mapping of short-read sequencing data (**Figure 3A**), we found that the read mapping rate aligned to the constructed pear pangenome was higher than that aligned to the PsbM reference genome (**Figure 3B**). The pear pangenome shows a further improvement in BUSCOs evaluation with a higher complete BUSCOs score (93.5%) and a lower missing (3.2%) and fragmented (3.3%) BUSCOs than the PsbM genome (complete BUSCOs: 92.9%; missing BUSCOs: 3.4% and fragmented BUSCOs: 3.4%) (**Table S11**). These findings demonstrate that compared with a single reference genome, the pear pangenome can reduce the reference bias and capture the missing genetic diversity. Moreover, pangenome modeling revealed a closed pan-genome, suggesting our constructed pear pangenome can capture nearly all the pear gene contents (**Figure 3C**).

**Figure 3.**
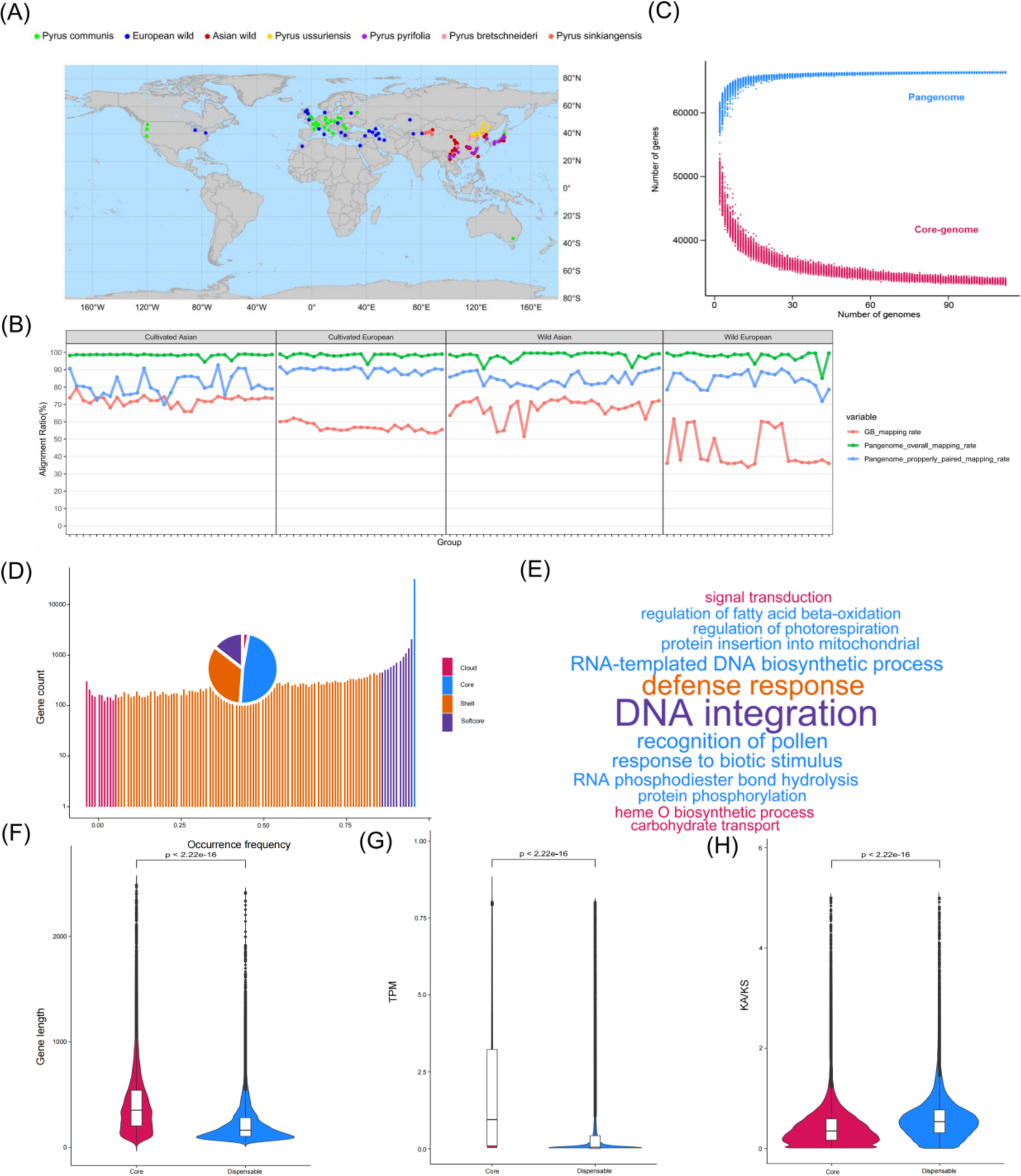
Pangenome of pear. (A) Geographical distribution of 113 pear resources. (B) Alignment ratio of 113 pear resources based on pangenome and single genome. (C) Pan-genome modeling. The pan-genome modeling shows no more dramatic increases when the number of accession genomes is over 30, indicating that the pear pangenome is sufficient to capture the majority of PAVs within pears. The upper and lower lines represent the pan-genome number and core-genome number, respectively. (D) Pangenome gene classification. (E) Word cloud of the GO enrichment of biological process for variable genes. (F) Boxplots show the median and upper and lower quartiles of gene length. (G) the RNA-seq expression level (H) Non-synonymous/synonymous substitution ratio (Ka/Ks) of core and dispensable orthologous gene clusters.

To detect the population-wide SNPs and PAVs in the pear pangenome, we further aligned 113 pear resequencing data (Wu et al., 2018) to the pear pangenome in which these pear accessions were from worldwide collections representing the genetic diversity of cultivated and wild pear species (**Figure 3A**). We identified 18,667,069 SNPs with 1,158,812 (5.4%) SNPs located in the non-reference sequences and further performed the SNP variant annotation to assess their functional effect. Subsequently, we found that SNPs located in non-reference sequences were more likely to have significant impacts on gene function, with 59.3% and 2.7% of the SNPs located in non-reference sequences and a lower percentage of SNPs (51.2% and 0.9%) located in reference sequences resulting in missense and nonsense mutations respectively (**Table S12**). In addition, a total of 326,461 PAVs ranging from 100 bp to 832.3 kb were identified across 113 diverse pear accessions using the pear pangenome. Based on the PAVs detection result, 32,174 (48.4%) were core genes and present in all 113 pear accessions, while 51.6% (34,367) of the genes were classified as dispensable, comprising 9,624 softcore, 22,885 shell, and 1,858 cloud genes, which were defined with a present frequency of more than 99%, 1-99%, and less than 1% respectively (**Figure 3D**). The proportion of dispensable genes observed in this study is relatively higher compared with other fruit plants such as apple (18.7%) (Sun et al., 2020), tomato (26%) (Gao et al., 2019), and watermelon (30.5%) (Wu et al., 2023). However, the proportion is similar to cucumber (43.8%) (Li et al., 2022) and smaller than citrus (64.6%) (Gao et al., 2023). Gene ontology (GO) enrichment analysis shows that genes with essential biological functions, including DNA-templated transcription, transmembrane transport and response to cadmium ion were enriched in core genes (**Table S13**). By contrast, genes with functions in pollen recognition, regulation of photorespiration, biotic stimulus response, and disease resistance are enriched in dispensable genes (**Figure 3E** and **Table S14**). Genes in dispensable orthologous clusters have shorter gene lengths, lower gene expression levels and higher non-synonymous/synonymous substitution ratio (Ka/Ks) (**Figure 3F-H**). In line with the cucumber pangenome study (Li et al., 2022), our results suggest dispensable genes are under stronger diversifying selection and a faster evolutionary rate than core genes and may be associated with phenological changes and disease resistance variations during selection for adaption to different environments.

### Characterization of the pear population using the pear pangenome

We further used our constructed pangenome to characterize the pear population. Calculation based on the SNPs makers, the nucleotide diversity (π) of pear at the pangenome level across all 113 pear accessions (Wu et al., 2018) was 4.84 x 10^-3^. Similar to the previous findings (Wu et al., 2018), the cultivated Asian and European pears have similar levels of nucleotide diversity (3.56 x 10^-3^) but lower than the wild Asian (4.23 x 10^-3^) and European pears (3.92 x 10^-3^) (**Figure 4A**). Using PAVs as markers, we found that the PAV diversity of cultivated (1.97 x 10^-3^) and wild (2.42 x 10^-3^) European pears was lower than cultivated (2.77 x 10^-3^) and wild (2.88 x 10^-3^) Asian pears (**Figure 4B**), suggesting that European pears contained fewer PAVs than the Asian pears at the pangenome level. In addition, using the pear pangenome, we further studied the population structure of 113 diverse pears. The cultivated and wild Asian and European pears can be separated into four clusters at both the SNPs based and PAVs based PCA (**Figure 4C and D, Figure S1 and S2**) and phylogeny results (**Figure 4E and F**). Interestingly, we found that PAVs based PCA can better distinguish the cultivated and wild European pears (**Figure 4D**). In the PAV-based phylogeny structure, wild Asian pears *P. calleryana*, *P. dimorphophylla,* and *P. phaeocarpa* show a closer phylogenetic relationship with cultivated Asian pear *P. pyrifolia* (**Figure F**), which is not reflected in the SNP-based phylogeny structure (**Figure D**). These results suggest that PAVs-based analysis at the pangenome level provides additional insight into the characterization of population structure in pears.

**Figure 4.**
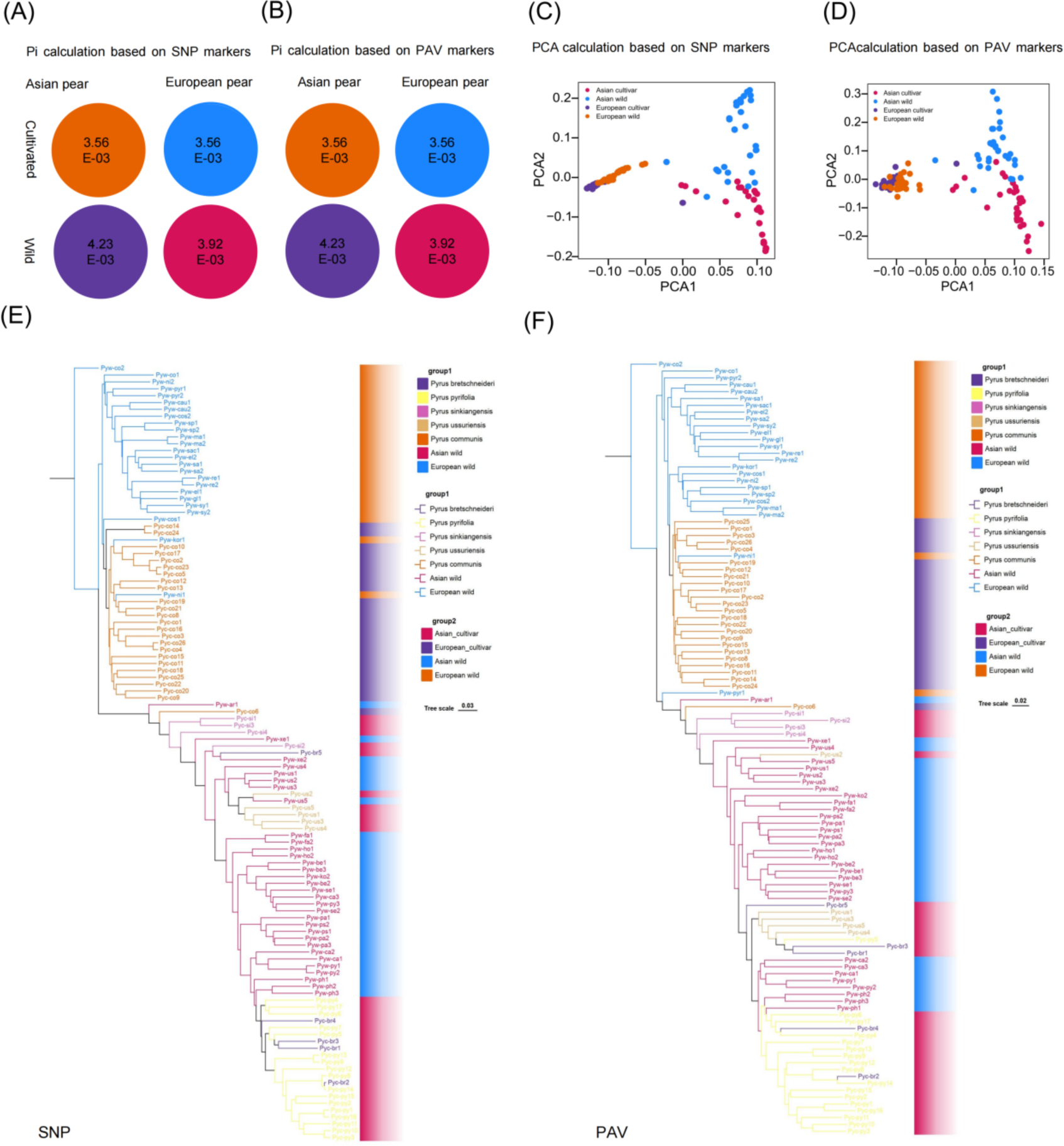
Characterization of the pear population basing on the pear pangenome. (A) and (B) represented the calculation of nucleotide diversity based on PAVs and SNPs. Nucleotide diversity (π) across 113 pear accessions at the pangenome level was 4.84 x 10^-3, with cultivated Asian and European pears showing similar levels (3.56 x 10^-3), which are lower than their wild species, while PAV diversity indicated European pears (both cultivated and wild) have lower diversity than Asian pears.(C) and (D) represented the PCA analysis based on SNPs and PAVs reveals different patterns in the pear population structure.(E) and (F) denoted the phylogenetic trees constructed based on SNPs and PAVs reveal different patterns in the pear population structure.

### Gene PAVs provide new insight into the selection footprints of European and Asian pears

Analyzing gene content across diverse pear accessions demonstrated a significant difference in average gene number per individual between European pear and Asian pear. European pear contains a higher average number of genes (56,842), with a reduction in Asian pear (56,118) (**Figure 5A**). Interestingly, different from previous findings that gene number reduction was observed during crop domestication and breeding improvement (Bayer et al., 2022; Gao et al., 2019), cultivated pears including cultivated European pear and cultivated Asian pear do not have significantly different gene numbers than wild European pear and wild Asian pear respectively (**Figure S3**). The reduction in average gene numbers hides a more complex pattern of increases and decreases in the frequency of specific genes across the population. To identify gene PAV changes across diverse pear accessions, we compared gene occurrence frequencies between European pear and Asian pear and between their wild and cultivated individuals (**Figure 5B**). A total of 8,361 genes showed a higher occurrence frequency in European pears (**Table S15**), while 8,093 genes showed an increased occurrence frequency in Asian pears (**Table S16**). GO enrichment analysis revealed that genes showing a higher occurrence frequency in Asian pears are associated with the function of defense response, response to biotic stimulus, vessel member cell differentiation, and sterol biosynthesis (**Table S17**).

**Figure 5.**
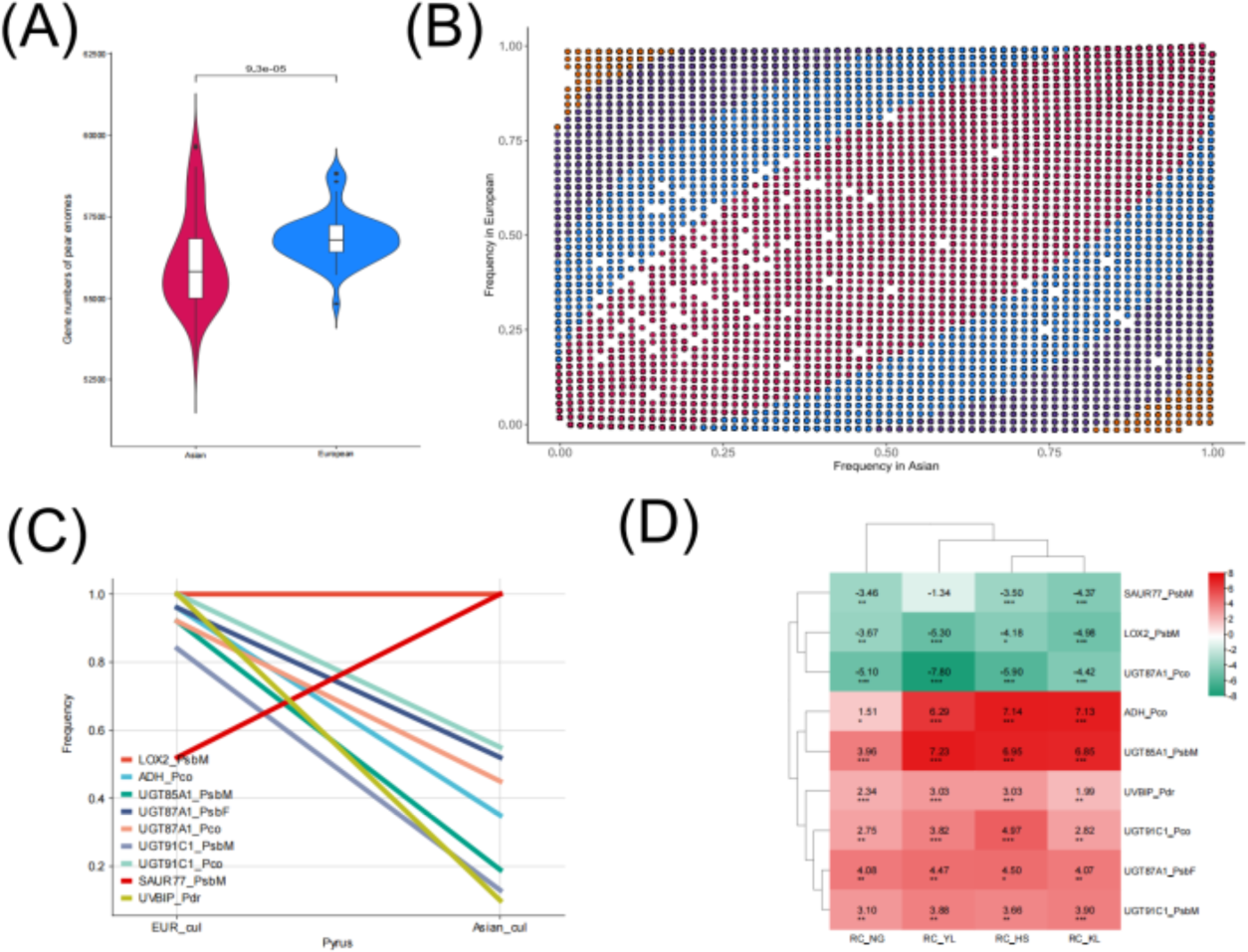
Gene PAVs between European and Asian pears. (A) The gene number between Asian and European pear genomes. (B) Change of gene PAV occurrence frequency between Asian and European pear genomes. (C) The change of PAV occurrence frequency of nine genes between Asian and European pear genomes. These nine genes may play key roles in fruit quality trait differentiation between Asian and European pear genomes. (D) The differentiation of gene expression among nine key genes between Asian and European pear genomes.

To further investigate how PAVs affect fruit quality traits in pears, we identified genes showing significant differences in PAV occurrence frequency between Asian and European pear populations that are functionally associated with fruit shape, aroma, color, stone cells, and fruit hardness. In total, we identified 192 fruit quality traits associated genes (**Table S18**), with homologous six genes exhibiting significant correlations with fruit shape development, including transcription repressor gene *OFP16* (*PsbM011G02493* and *PsbM005G03117*) (Snouffer et al., 2020), homeobox protein knotted-1-like gene *KNOX1* (*Pco_Presence_Seq_006877*) (Wang et al., 2022), *SUN domain-containing protein 5 gene* (*Pdr_Presence_Seq_013651*) (Ma et al., 2022), and *TRM (PsbM003G00721* and *Pco_Presence_Seq_003059*) (Zhang et al., 2023). For example, the gene *PsbM005G03117* is present in all Asian pear accessions but less than half of European pear accessions (Occurency frequency: 0.55) (Differential occurrence frequency: 0.45, P-value: 1.54E-08), while *PsbM011G02493* showing a different pattern than present in all European pear accessions and near three-quarters of Asian pear accessions (Occurrency frequency: 0.72) (Differential occurrence frequency: 0.28, P-value: 1.06E-05). Interestingly, the largest differential occurrence frequency (0.67) between Asian and European pears is observed in the gene *SUN5* (*Pdr_Presence_Seq_013651*) (Occurrence frequency in Asian and European pear: 0.33 and 1, P-value: 1.56E-14). Besides, we identified 28 genes associated with fruit color development, including 12 LOB domain-containing protein (*LBD*) genes, 10 B-box zinc finger protein (*BBX*) genes (Bu et al., 2022) and 4 anthocyanin regulatory (Myb_DNA-binding) protein genes (Jin et al., 2016). Among these genes, a LOB domain-containing protein *CRL* gene (*PsbM001G01311*) shows a significantly lower occurrence frequency in European pears compared with Asian pears, especially cultivated European pear (Differential occurrence frequency between cultivated Asian and European pears: 0.88, P-value: 5.00E-11), while a B-box zinc finger protein 22 gene (*Pdr_Presence_Seq_002427*) is present in all cultivated European pear but nearly absent in all cultivated Asian pear (Differential occurrence frequency between cultivated Asian and European pears: 0.96, P-value: 8.40E-13). Moreover, we identified 57, 3, and 102 genes playing important roles in the development of fruit stone cells, fruit aroma, and firmness show differences in PAV occurrence frequency. Additionally, we also observed that the UV-B-induced protein gene *UVBIP* (*Pdr_Presence_Seq_004483*) (Qian et al., 2014) regulating photomorphogenic responses to UV-B and blue light shows higher occurrence frequency in European pears (Differential occurrence frequency between cultivated Asian and European pears: 0.92, P-value: 2.54E-24) (**Figure 5C**).

We further performed gene expression analysis merging RNA-seq data from five pear species and pear fruit tissues of seven different stages (Zhang et al., 2016). Subsequently, we identified six fruit firmness and two aroma-related genes showing significant PAV occurrence frequency as well as significant differential gene expression between Asian and European pear populations. Notably, compared with the Asian pears, the fruit hardness development genes including the UDP-glycosyltransferase gene *UGT87A1* (Wu et al., 2017) (*Pco_Presence_Seq_002869*, Differential occurrence frequency between cultivated Asian and European pears: 0.43, P-value: 1.75E-05) and the auxin-responsive protein gene *SAUR* (Zhang et al., 2022) (*PsbM005G02020*, Differential occurrence frequency between cultivated Asian and European pears: −0.34, P-value: 5.76E-07) show a significant downregulation in expression in European pear fruit tissues (log2FoldChange >3.46). However, the other four UDP-glycosyltransferase genes (*Pco_Presence_Seq_001484*, *PsbF_Presence_Seq_007967*, *PsbM015G01882* and *PsbM017G01217*) having a higher occurrence frequency in European pears (Differential occurrence frequency between cultivated Asian and European pears >0.22, P-value< 5.26E-04), showed an upregulated expression in European pears (log2FoldChange >2). Among these genes, the gene *PsbF_Presence_Seq_007967*, a homolog of *Pbr011574*, was reported to be located in the fruit firmness QTL according to a previous study. In addition, two aroma-related genes alcohol dehydrogenase (*ADH*) (Shi et al., 2019; Zhang et al., 2010) and Lipoxygenase 2 gene (*LOX*) (Luo et al., 2021) also show significant differential gene expression between Asian and European pear populations. For example, the *ADH* gene (*Pco_Presence_Seq_011843*) displays a higher occurrence frequency (Differential occurrence frequency between cultivated Asian and European pears: 0.45, P-value: 3.17E-06) as well as an upregulated expression in European pears (log2FoldChange>1.51). Furthermore, we observed the *UVBIP* gene (*Pdr_Presence_Seq_004483*) with a higher occurrence frequency in European pears, shows a significantly elevated expression in European pears compared to Asian pears (**Figure 5D**). The above results suggest that gene PAVs might play key roles in differentiating pear quality traits between Asian and European pears by regulating the expression of related genes.

### Exploring Structural Variation, Expression, and Functional Validation of the Sepal Abscission Zone Pear Gene *GH3.1*

Based on the previous studies (Guo et al., 2022; Qi et al., 2013), *GH3.1* (*PsbM002G01431*), encodes an indole-3-acetate synthetase which is a pear sepal development candidate gene and plays an important role in early auxin response Analysis of allele-specific structural variations within the gene *GH3.1* promoter unveiled a 104bp insertion (SV104) in the genome of PsbM compared to the genome of PsbF, Pbr and Ppy, spanning from position 11,287,633 to 11,407,583 on the upstream promoter of *GH3.1* (**Figure 6A**). In addition, an SNP1 (G→ A) in the first intron of the gene *GH3* potentially affecting gene alternative splicing was identified in PsbM and Pbe (Mature fruit with completely shedding of sepals). We conducted a further examination of the distribution of both the insertion and SNP in a diverse collection of 113 pear accessions. Intriguingly, we identified a total of five accessions (comprising 4 from cultivated and wild Asian pears, namely Pyc_br1, Pyc_br5, Pyw_us5, Pyw_xe1, and 1 from wild European pear Pyw_cau1) (**Table S19**) that harbored the same SNP as PsbM, while an additional six accessions (including four from Asian cultivated pears Pyc_py5, Pyc_py7, Pyc_si1, Pyc_us1, and two from wild pears Pyw_pa1 and Pyw_pa3) (**Table S20**) exhibited the insertion. Furthermore, the expression of the *PsbMGH3.1* gene in the sepal abscission zone of the paternal haplotype genome PsbM (‘Xuehuali’) was five times higher than that in the maternal haplotype genome PsbF (‘Kuerlexiangli’, mature fruit with incompletely shedding of sepals) (**Figure 6B** and **Table S21**). Notably, these shared genetic variants aligned precisely with the paternal genotype of the ‘Yuluxiang’ pear haploid genome, providing insights into the genetic diversity of this distinct lineage.

**Figure 6.**
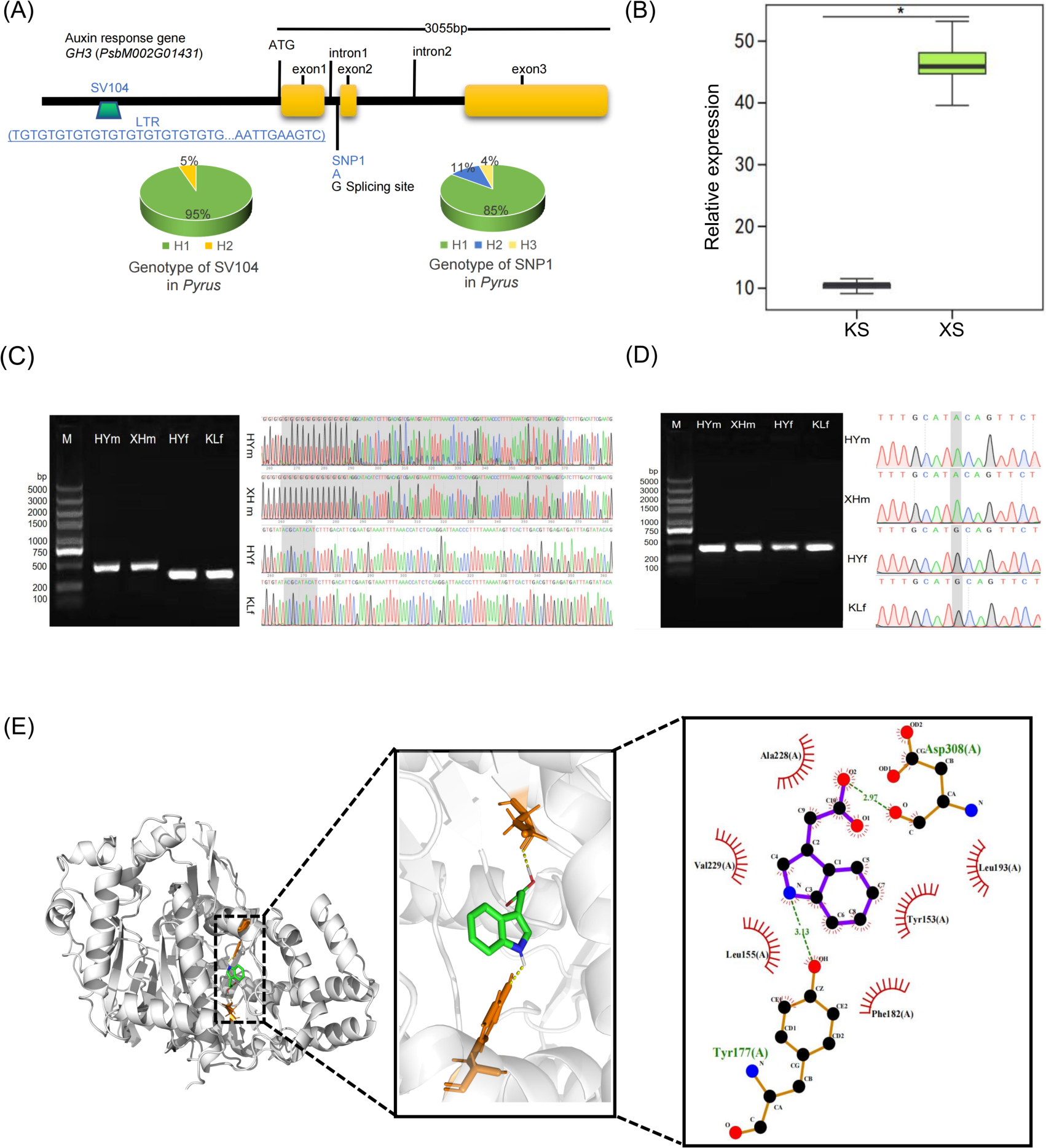
The characterization of the sepal abscission zone pear gene GH3.1 from structural variation, expression to functional validation. (A) The variation of gene *GH3* in Pyrus. (B) XS and KS represent the late stage of sepal development of ‘Xuehuali’ and ‘Kuerlexiangli’ respectively, n=3,* P < 0.05. (C) The left part: The inserted mutation sites of ‘Yuluxiang’ pear and its parents, M: Marker, HYm and MH, and HYf and FK, represent the cloning fragments of the allele *PsbMGH3.1* from upstream to downstream of the mutation sites of ‘Yuluxiangli’ and ‘Xuehuali’, respectively, and the cloning fragments of the allele *PsbFGH3.1* from upstream to downstream of ‘Yuluxiang’ pear and ‘Kuerlexiangli’; The right part: Detection of genotyping of mutation sites. (D) The left part: The mutated sites of ‘Yuluxiang’ pear and its parents. (E) The left part: Simulation of molecular docking between PsbMGH3.1 and IAA, and local amplification of active pocket; The middle part: IAA chemical structure formula; The right part: Hydrogen bond force between PsbMGH3.1 and IAA.

We further designed primers based on the flanking sequences of the insertion for validation, confirming the precise alignment of cloned variant sites in the ‘Yuluxiang’ pear genome and its parental genome with those in the assembled doubled haploid genomes (PsbF and PsbM) (**Figure 6C and D**). Utilizing the protein crystal structure of *VvGH3.1* (Peat et al., 2012; Wang et al., 2015) from grape (PDB ID: 4B2G) and IAA (PubChem ID: 802), we constructed a molecular docking model, revealing the active pocket and binding site predictions of PsbMGH3.1 with IAA. Among nine identified binding modes, the final docking conformation with an optimal binding energy of −7.0 kcal/mol was selected (**Table S22** and **Figure 6E**). The interactions between IAA and PsbMGH3.1 were primarily characterized by hydrogen bonding with Asp308 and Tyr177, with spatial distances of 2.97 Å and 3.13 Å, respectively. Notably, overexpression of the sepal shedding gene PsbMGH3.1 in rice led to noticeable alterations in transgenic plant phenotypes, including significantly reduced adventitious root length, longest leaf length, and plant height compared to wild plants (**Table S23**, **Figure S4, and Figure S5**).

## Discussion

In this study, we *de novo* assembled and annotated the genomes of the maternal (PsbM) and paternal (PsbF) lines of the hybrid ‘Yuluxiang’ pear using the PacBio long read and Hi-C sequencing. Compared with other published pear genomes, our pear genome assemblies show significant completeness and accuracy, allowing the characterization of complex centromere and telomere regions in nearly all chromosomes. Our high-quality genome assemblies also enable a comprehensive structural variation discovery among genomes of different pear species, leading to the identification of 6,341 high-confident SVs (7 inversions, 15 translocations, 3,989 deletions, and 2,330 insertions). Along with previous studies of *Brassicaceae* (Dolatabadian et al., 2022) and soybean (Bayer et al., 2021), we found that deletions detected in Pco, Pdr and PsbF genomes result in a complete loss of two copies of disease-resistance related genes, reflecting the differentiation of disease resistance in pear species. Hence, these two high-quality reference genomes provide valuable genomic resources for detecting genomic variations in diverse pear species.

To capture the complete genetic diversity of pears, we constructed the first pear pangenome using the PsbM reference genome as the backbone genome and identified 21,224 novel genes in the non-reference genome. By aligning the read sequencing data to the PsbM reference and the pear pangenome, we identified a higher mapping rate, and a larger number of SNPs in the pear pangenome, reflecting that the pear pangenome can effectively reduce the single reference bias for mapping and variant calling. In contrast to SNP markers, our analysis based on PAVs provides additional insights into the population structure of pears, which is similar to previous findings in *Arabidopsis* (Jiao and Schneeberger, 2020), *Amborella* (Hu et al., 2022), and rice (Wang et al., 2023a). Furthermore, based on the pangenome, we first report distinct PAV occurrence frequencies and gene expression patterns in Asian and European populations for genes linked to pear fruit quality traits, including fruit shape, stone cells, aroma and fruit firmness. For instance, The *ADH* gene (Shi et al., 2019; Zhang et al., 2010) (*Pco_Presence_Seq_011843*) exhibits a higher frequent occurrence in European populations, alongside upregulated expression in European pears. These results are consistent with previous findings that SVs have significant impacts on fruit quality traits such as shape, flesh color, and sucrose content (Song et al., 2020; Zhou et al., 2019; Lyu et al., 2023). While the association between the PAV gene and gene expression levels remains unclear, additional research is required to elucidate the regulatory mechanisms of how PAV influences gene expression.

Using the constructed pear pangenome, a 104bp structural variation upstream of the *PsbMGH3.1* gene promoter was identified, contrasting with the corresponding position in *PsbFGH3.1*. This variation has the potential to augment gene expression responsible for sepal abscission, potentially leading to increased sepal shedding in ‘Xuehuali’. Furthermore, we investigated the functional role of PsbMGH 3.1, an auxin nicotinamide synthase gene, recognized as an early responder in the indole-3-acetic acid (IAA) signaling pathway, critical in plant organ shedding encompassing biosynthesis, transportation, and catabolism processes (El Houari et al., 2023; Guo et al., 2022; Pencik et al., 2013; Qi et al., 2013). Additionally, IAA’s binding to amino acids such as IAA-leucine, IAA-alanine, and IAA-phenylalanine through amide bonds, as well as its storage in a bound state with amino acids, indirectly reducing intracellular free IAA content, is highlighted. Our study predicted potential hydrogen bond formation between indole-3-acetic acid (IAA) and Tyr177 and Asp308 of *PsbMGH3.1*, indicating the likelihood of *PsbGH3.1* interaction with IAA. Notably, *PsbMGH3.1* displayed upregulation in the detached zone of the paternal haplotype genomic cultivar ‘Xuhuali’, leading to a characteristic low-concentration auxin phenotype in heterologously overexpressing transgenic plants. We propose that *PsbMGH3.1* potentially influences pear sepal formation by modulating IAA content in plant cells, a critical aspect given the impact of persistent sepals on pear fruit quality and the significance of understanding sepal shedding at the molecular level for the cultivation of sepal-shedding fruits.

## Materials and Methods

### Materials

#### Plant sequencing materials

The prevalent cultivar ‘Yuluxiang’ (*P. sinkiangensis ×P. bretschneideri*) (Ding et al., 2021) grown in the Fruit Tree Research Institute of Shanxi Agricultural University (N37.21 °, E112.30 °) of China was used for doubled haploid de novo sequencing. The collected leaves were used for HiFi sequencing and HiC fixation and library construction, the collected leaves of ‘Yuluxiang’ parent were used for Illumina sequencing to assist genome assembly, and the mixed collected tissues of leaves, flowers (including pistils and stamens), sepals, stems, and young fruits were used for ONT library construction to assist in predicting the genes of ‘Yuluxiang’ pear. Two weeks after the flowering period, sepal abscission zone from the parents of ‘Yuluxiang’ pear (‘Kuerlexiangli’ and ‘Xuehuali’) were collected for cloning the target fragment of GH3.1 gene and fluorescence quantitative PCR.

### Methods

#### DNA and RNA extraction

Total genomic DNA and RNA were extracted from the sepals of the abscission zone of ‘Yuluxiang’ pear and its parent plants, using the CTAB method. Extracted nucleic acids were then quantified using a Nanodrop spectrophotometer and assessed for fragment integrity using an Agilent 2100 Bioanalyzer, to ensure their suitability for subsequent genotype cloning (**Table S24**).

For ‘Kuerlexiangli’ and ‘Xuehuali’, RNA was isolated from the sepals of the abscission zone at the end of their developmental stage using the CTAB method. This RNA was then reverse transcribed into double-stranded cDNA. The cDNA was used in quantitative fluorescence PCR according to the previously published research (Ding et al., 2021) (**Table S25**) and for gene cloning procedures (**Table S 26**).

#### Illumina, ONT, PacBio and Hi-C library construction and sequencing

The qualified 350 bp fragment of DNA was selected to perform pair ended sequencing according to the Illumina library of according to the Illumina HiSeq X ten (150bp) system standard process (Wright et al., 2017). Qualified RNA from various tissues was extracted and mixed equally for ONT library. After breaking the RNA into an average of 8 kb fragments for library construction and ONT sequencing based on the structural differences of nucleotide sequences (Leger et al., 2021). The genomic DNA was broken into 15 kb fragments by the Megaruptor ® 2 and the SMRT bell library was constructed using SMRT bell Express Template Prep kit 2.0 (Pacific Biosciences), 10 µg of DNA was taken to perform SMRT sequencing on the Sequel II system. After correcting the sequencing error high fidelity HiFi reads were obtained. More than 2 g of fresh and completely natural condition of growing ‘Yuluxiang’ pear’s young branches (cleaned with distilled water in advance) was collected and qualified for Hi-C library. DNA products was obtained through fragmentation, fixation, cell lysis, and enzyme digestion, end labeling, flat end connection, purification, etc. Afterwards, 2 µg of DNA samples were taken and interrupted using the Covaris S 220 interrupter. According to the BMK plant Hi-C SOP (BMK190227, Biomarker Technologies, Beijing) reaction system, the DNA ends were repaired, fragments from 300 bp to 700 bp were selected and subjected to quality testing (Qubit3.0 and qPCR were used to detect fragment integrity and concentration). After constructing the library, sequencing was performed using the ilumina platform (PE150).

#### De novo genome assembly and Hi-C scaffolding

Based on hifiasm software (Conesa et al., 2005) and incorporating the resequencing data from ‘Yuluxiang’ pear parents, kmer files for both parent strains were generated. This was achieved using the yak software, with the command hifiasm -o hifiasm.asm -t 64 −1 R01.yak −2 R02.yak ccs.fastq.gz. Subsequently, the resulting data were bifurcated into two distinct haplotype genomes: PsbM and PsbF. The subsequent genome scaffolding employed the LACHESIS software (Burton et al., 2013) to facilitate the grouping, ordering, and orientation of genomic sequences. Manual mapping and meticulous scrutiny were then performed, culminating in the acquisition of a chromosome-level, doubled genome.

#### Repetitive sequence prediction

The assembled ‘Yuluxiang’ pear’s dual haplotype genome sequences were initially processed using LTR_Harves (Flynn et al., 2020). Subsequently, the LTR_Finder program (Ellinghaus et al., 2008) was employed for the prediction of long terminal repeat (LTR) sequences. The outputs from the aforementioned software were consolidated using LTR_Retriever (Ou and Jiang, 2018), employing a neutral mutation rate of u=1.3e-8 per base pair per year. Following this, both haplotype genome sequences were subjected to Misa.pl (Ou and Jiang, 2018) and the trf program (Tarailo-Graovac and Chen, 2009). The objective was to forecast microsatellite sequences (SSRs) and tandem repeat sequences (Benson, 1999). This process culminated in the generation of an integrated repetitive sequence library file for the dual haplotype genome. Finally, the RepeatMasker software (Flynn et al., 2020) utilized the library file, invoking the -lib parameter. This produced the ultimate repetitive sequence, wherein repetitive sequence regions were substituted with ‘N’. Post this, repetitive sequences were excised, streamlining the prediction of the doubled haplotype genes.

#### Gene annotation

Extract high-quality full-length transcripts of ‘Yuluxiang’ pear were used for gene prediction model training based on Nanopore full-length transcriptome data of mixed tissues (leaves, flowers, sepals, stems, fruits, etc.). The raw sequencing data of Nanopore were filtered and obtained by Pychopper (https://github.com/friend0/pyChopper), then the filtered data were treated by the Pinfish (https://github.com/nanoporetech/pipeline-pinfish-analysis)package, Transdecoder (Perina et al., 2017), maker2 (Holt and Yandell, 2011), AUGUSTUS (Stanke et al., 2006) to train the gene prediction model of ‘Yuluxiang’ pear, and produced a standard .gff format file by EVM (Haas et al., 2008). Finally, the optimal alignment of genes function were inferred using diamond (Buchfink et al., 2015) against KEGG database (Kanehisa et al., 2012), E=1E-5, NCBI non redundant (NR) database (Pruitt et al., 2007), EggNOG (Huerta-Cepas et al., 2019), TrEMBL [205] (O’Donovan et al., 2002), InterPro (Mitchell et al., 2015), and SwissProt (Boeckmann et al., 2003) protein databases.

#### Non coding RNA prediction

tRNA and rRNA sequences were annotated utilizing tRNAscan SE (Lowe and Eddy, 1997) and the barrnap algorithm (v0.9) and both were executed with default parameters. For the identification of miRNAs and snRNAs, the PRfam database (Finn et al., 2014) (v14.5) was employed, leveraging the Infernal software (Nawrocki and Eddy, 2013) (v1.1.1) under default settings.

#### Structural variations detection

We used the PsbM genome as the reference for SV detection. A European wild pear, a Chinese wild pear, a Xinjiang Pear (PbsF), a Japanese pear, a Chinese white pear and a dwarfing pear were aligned to the PsbM genome, respectively, using Mummer (Marcais et al., 2018) v4.0 with the parameters (-l 50 -c 100 -maxmatch). The initial alignment results were filtered using delta-filter with parameters (-m -i 90 -l 100). The resulting filtered delta files were used as the input for the SyRI pipeline with default parameters (Goel et al., 2019). The detected variations from SyRI comprise genome rearrangements and sequential variations, which can occur in both rearrangement and syntenic regions. Therefore, the sequential variants embedded in the rearrangements were not included for further analysis. According to the definitions of sequence variation in SyRI outputs, we converted these variations into four types of SVs: Insertion (INS and CPG variants), Deletion (DEL and CPL variants), Inversion (INV and INVDP variants), and Translocation (TRANS variants). SyRI results were visualized using the plotsr package. SURVIVOR (Jeffares et al., 2017) was used to merge (with parameters: 1000 2 1 1 0 50) the individual genotyped VCF files.

#### Pangenome construction

The pear pangenome was constructed using a similar method described in Song et al.’s (2020) and Tahir ul Qamar M. et al.’s (2019) studies. In detail, the PAV sequences in the **other seven pear genomes** relative to the PsbM reference genome were identified using the Mummer ‘show-diff’ function (Marcais et al., 2018). Initially, sequences intersecting with gap regions within the respective genome were excluded from consideration. Additionally, sequences aligning with gap-start or gap-end boundaries were filtered out. To pinpoint unique sequences within each genome, candidate PAV sequences were aligned to the PsbM genome using minimap2 (Li, 2018) with the parameter setting ‘-x asm10’. Redundant sequences (PAV sequences showing larger than 80% similarity and coverage with the PsbM genome) were filtered out. The rest of the PAV sequences were retained and linked with 100 Ns bases as the final PAV regions. Genes with at least 80% coding sequence region overlapping with PAV regions were designated as novel genes.

#### SNPs calling

Clean reads of 113 diverse pear accessions (Wu et al., 2018) were mapped to the constructed pear pangenome using BWA MEM (Li, 2013) v0.7.17. Default settings were used and duplicates were removed by Picard tools (http://broadinstitute.github.io/picard/). Reads were realigned by GATK v3.8-1-0 RealignerTargetCreator and IndelRealigner (McKenna et al., 2010), followed by variant calling using GATK HaplotypeCaller. The resulting SNPs were filtered (QD < 2.0 || MQ < 40.0 || FS > 60.0 || QUAL < 60.0 || MQrankSum < −12.5 || Read-PosRankSum < −8.0) to remove low-quality SNPs. High-confidence SNPs were obtained by further filtering out the SNPs with minor allele frequency <0.05 and missing genotype rate > 10% using VCFtools (Danecek et al., 2011).

#### Gene PAV analysis

Short-read sequencing data of 113 diverse pear accessions were aligned to the constructed pear pangenome using Bowtie2 v2.3.3.1 (Langmead and Salzberg, 2012) (–end-to-end –sensitive -I 0 -X 1000). We used the gene PAV detection approach described in Hu et al.’s study (2020). A gene was classified as absent when the horizontal coverage across exons of the gene was <20% and the vertical coverage was <2×. A gene PAV matrix was generated using this threshold, with each gene classified as presence or absence for each accession. Statistical significance of PAV frequency changes between European and Asian pear populations was determined by Fisher’s exact test. P-values were adjusted for multiple comparisons using the Bonferroni method. PAVs with an adjusted p-value < 0.01 and differential occurrence frequency between groups ≥0.2 were identified as under significant occurrence frequency changes between populations.

#### Population analysis

The SNP and PAV genotype matrices were used to perform the SNP and PAV based population analysis. Nucleotide diversity values (π) were calculated using Pixy (Korunes and Samuk, 2021) using a 100-kb window. The phylogenetic trees of 113 diverse pear accessions were constructed using IQ-tree using a maximum likelihood method (with the parameters -alrt 1000 -bb 1000) and visualized using ITOL (Letunic and Bork, 2019). Principal component analysis (PCA) was performed using the gcta (Yang et al., 2011) v1.94.

#### GO analysis

Gene function annotation was performed using Blast2GO (Conesa et al., 2005) v2.5. Genes in the pangenome were aligned to the proteins in the Viridiplantae database using BLASTP (Camacho et al., 2009) (E-values <1 × 10-5). Gene ontology (GO) analysis was conducted using topGO (Raudvere et al., 2019) and Fisher’s exact test with ‘elim’ used to correct for multiple comparisons.

#### RNA-seq differential expression analysis

Clean RNA sequencing reads (Zhang et al., 2016) were generated by removing low-quality reads, adapters, and reads containing poly-N using the fastp (Chen et al., 2018) v0.20 with default parameters. Clean reads were further mapped to the constructed pear pangenome using Hisat2 (Pertea et al., 2016) v2.1.0 with default parameters, and the number of mapped reads was counted using HTSeq (Anders et al., 2015) v0.6.1. RNA-Seq reads were normalized to TPM (Transcripts Per Kilobase Million) and low expression reads (TPM < 1) were removed. Differential expression analysis was performed using the DESeq2 (Love et al., 2014) R package 1.18.0 between the European pear population (RC: Red Clapp’s Favorite mutant, *P. communis*) and the Asian pear populations (HS: Hosui, *P. pyrifolia*; YL: Yali, *P. bretschneideri*; KRLXL: Kuerlexiangli, *P. sinkiangensis*; NG: Nanguo, *P. ussuriensis*).

#### Functional validation of the heterologous overexpression

Sterilization was carried out on rice seeds at the grain-filling stage using the following steps. Seeds with full grains were selected and then rinsed thoroughly with clean water. The selected seeds were placed in a beaker and sterilized using 70% alcohol for two minutes. Subsequently, a 30% NaClO solution was added until the beaker was filled, and the seeds were left in the solution for 90 minutes. After discarding the NaClO solution, the sterilized seeds were rinsed at least four times with deionized water (ddH2O) to remove any remaining surface residues

#### Gene cloning, quantification, and transgenic materials

The abscission zone tissues of the pear variety ‘Yuluxiang’ pear and its parents of ‘Kuerlexiangli’ (*Pyrus sinkiangensis*, represented maternal haplotype genome PsbF) and ‘Xuehuali’ (*Pyrus bretschneideri*, represented paternal haplotype genome PsbM) were collected at the end of sepal development (two weeks after blooming) in the pear resource garden (Taigu) of the Fruit Tree Research Institute of Shanxi Agricultural University (E112°48’ N37°38’). Seeds at the grain-filling stage were used in rice (*Oryza sativa* L. cv. Nipponbare).

## Methods

### Construction of overexpressed vector of the pEGOEP35S-H-PsbMGH3.1-GFP

According to the designed cloning primers, after PCR amplification of target fragments, ligation-mediated restriction, recombinant plasmid transformation, and colony plasmid extraction, the quality of the extracted plasmids was measured by agarose gel electrophoresis at a concentration of 1% (Figure S6 and S7). Finally, Sanger sequencing was performed based on the primer eGFP-cx, and the target genes were aligned with the de novo assembled doubled haploid genomes.

### Callus induction and Subculture

Using sterilized tweezers, the pre-treated seeds were handled carefully. Young rice embryos were then isolated using sterilized steel hooks and placed on sterile filter paper. These embryos were transferred onto an induction culture medium. Afterward, the Petri dish was sealed with plastic wrap and incubated under dark conditions at 25 ℃ for five days. After that the embryogenic callus tissues were selected for further subculturing over a three-day period. Subsequently, the constructed Agrobacterium vectors were co-cultured with the induced tissue and subjected to antibiotic screening, induced differentiation, and rooting steps to obtain a preliminary positive transgenic seedling (Tie et al., 2012).

### Molecular docking

The crystal structure of the VvGH3.1 protein from grape (PDB ID: 4B2G) was obtained from the Protein Data Bank (PDB) database (https://www.rcsb.org/). Similarly, the chemical structure of IAA (PubChem ID: 802) was downloaded from the PubChem database. These structures were then used in docking studies conducted using Autodock (Li et al., 2022). The study involved docking IAA with the protein encoded by the gene PsbMGH3.1 (Peat et al., 2012). Among the various conformations generated during docking, the one demonstrating the highest binding affinity was selected as the final docking conformation.

## Authors’ contributions

BD, HH, RX, LL, RV, and XG conceived and designed the experiments. BD, HH, and TL drafted the manuscript with contributions from RV, UlT, YL, YH, JQ, and JZ. BD, SY, XF, and HG managed sample collections, while BD, HH, SH, GH, and ZC conducted data analysis and performed statistical analyses. BD and TL executed the experiments. All authors have read and approved the final manuscript.

## Availability of data and materials

The raw genome sequenced data pertinent to this research have been deposited in the National Genomics Data Center, China National Center for Bioinformation / Beijing Institute of Genomics, Chinese Academy of Sciences (GSA: PRJCA018905 and CRA012175) and are publicly accessible at NGDC. The reported resequencing of 113 pear accessions from worldwide and RNA sequencing of pear fruit in five cultivars are detailed in Wu et al., (2018) and Zhang et al., 2018 in PRJNA381668 and PRJNA309745, separately. Yuluxiang pear double haploid genome and pear pan-genomes annotation can be accessed at 10.6084/m9.figshare.23950353.

## Acknowledgement

We extend our gratitude to Engineer Tao Huang from Beijing Baimaike Biotechnology Co., Ltd., for his invaluable assistance with genome sequencing and assembly, and to Dr. Renjian Li from the College of Plant Science & Technology, Huazhong Agricultural University, Wuhan, China, for his support with molecular docking.

## Supporting information

Additional supporting information might be avaliable online in the Supporting Information section at the end of the article.

Table S1 Chromosome-length assemblies of pear genomes (PsbF and PsbM).

Table S2 Statistics of centromere and telomere-specific repeats in both the PsbF and PsbM genomes.

Table S3 Statistics of repeat sequences within the assembled PsbF and PsbM genomes.

Table S4 Completeness of the PsbF and PsbM genomes.

Table S5 Functionally annotated gene number of PsbF and PsbM.

Table S6 Statistics of non coding RNA in PsbF and PsbM.

Table S7 Statistics of Large genomic variations in pear genomes.

Table S8 Statistics of high-confident SVs across seven pears genomes.

Table S9 Statistics of high-confident SVs in the pear genomes.

Table S10 Statistics of repetitive elements accounted for the non-reference genome than the reference genome (54.3%).

Table S11 Evaluation of BUSCOs in the pear pangenome.

Table S12 The distributed ratio of SNPs between reference sequences and non-reference sequences.

Table S13 Gene ontology (GO) enrichment analysis in core genes.

Table S14 Gene ontology (GO) enrichment analysis in dispensable genes.

Table S15 Characterization of higher occurrence frequency in European pears.

Table S16 Characterization of higher occurrence frequency in Asian pears.

Table S17 GO enrichment analysis showing a higher occurrence frequency in Asian pears.

Table S18 Identification of fruit quality trait related genes in PAV occurrence frequency between Asian and European pear populations.

Table S19 Genotype of SNP1 (GH3.1) in *Pyrus*.

Table S20 Genotype of SV1041 (GH3.1) in *Pyrus*.

Table S21 Expression of *PsbMGH3.1* in exfoliated calyx of tissues at the end of persistent abscission zone.

Table S22 Molecular binding energy of PsbM GH3.1 and IAA.

Table S23 Phenotype of transgenic *PsbMGH3.1* in *Nipponbare*.

Table S24 Genotyping sequence of targeted genes and theirs primers.

Table S25 Quantitative PCR primer information.

Table S26 Primer of sequence of CDS-cloning gene PsbMGH3.1.

Figure S1 SNPs based on PCA analysis in the pear population structure.

Figure S2 PAVs based on PCA analysis in the pear population structure.

Figure S3 Statistic of gene numbers in the pear population structure.

Figure S4 Phenotype of transgenic PsbMGH3.1 in rice (Oryza sativa ssp. japonica var. Nipponbare).

Figure S5 Statistics of phenotype of transgenic PsbMGH3.1 in rice (Oryza sativa ssp. japonica var. Nipponbare).

Figure S6 Electrophoresis of constructed map of transgenic vector.

Figure S7 Map of overexpression vector for pEGOEP35S-H-PsbM GH3-GFP.

## References

Anders, S., Pyl, P.T. and Huber, W. (2015) HTSeq--a Python framework to work with high-throughput sequencing data. Bioinformatics 31, 166–169.

Bayer, P.E., Hu, H., Petereit, J., Derbyshire, M.C., Varshney, R.K., Valliyodan, B., Nguyen, H.T., Batley, J. and Edwards, D. (2021) Yield is negatively correlated with nucleotide-binding leucine-rich repeat gene content in soybean. bioRxiv, 2021.2012. 2012.472330.

Bayer, P.E., Valliyodan, B., Hu, H., Marsh, J.I., Yuan, Y., Vuong, T.D., Patil, G., Song, Q., Batley, J., Varshney, R.K., Lam, H.M., Edwards, D. and Nguyen, H.T. (2022) Sequencing the USDA core soybean collection reveals gene loss during domestication and breeding. Plant Genome 15, e20109.

Benson, G. (1999) Tandem repeats finder: A program to analyze DNA sequences. Nucleic Acids Res 27, 573–580.

Boeckmann, B., Bairoch, A., Apweiler, R., Blatter, M.C., Estreicher, A., Gasteiger, E., Martin, M.J., Michoud, K., O’Donovan, C., Phan, I., Pilbout, S. and Schneider, M. (2003) The SWISS-PROT protein knowledgebase and its supplement TrEMBL in 2003. Nucleic Acids Res 31, 365–370.

Bu, Y.F., Wang, S., Li, C.Z., Fang, Y., Zhang, Y., Li, Q.Y., Wang, H.B., Chen, X.S. and Feng, S.Q. (2022) Transcriptome analysis of apples in high-temperature treatments reveals a role of MdLBD37 in the inhibition of anthocyanin accumulation. Int J Mol Sci 23, 3766.

Buchfink, B., Xie, C. and Huson, D.H. (2015) Fast and sensitive protein alignment using DIAMOND. Nat Methods 12, 59–60.

Burton, J.N., Adey, A., Patwardhan, R.P., Qiu, R., Kitzman, J.O. and Shendure, J. (2013) Chromosome-scale scaffolding of de novo genome assemblies based on chromatin interactions. Nat Biotechnol 31, 1119–1125.

Camacho, C., Coulouris, G., Avagyan, V., Ma, N., Papadopoulos, J., Bealer, K. and Madden, T.L. (2009) BLAST+: Architecture and applications. BMC Bioinformatics 10, 421.

Chagne, D., Crowhurst, R.N., Pindo, M., Thrimawithana, A., Deng, C., Ireland, H., Fiers, M., Dzierzon, H., Cestaro, A., Fontana, P., Bianco, L., Lu, A., Storey, R., Knabel, M., Saeed, M., Montanari, S., Kim, Y.K., Nicolini, D., Larger, S., Stefani, E., Allan, A.C., Bowen, J., Harvey, I., Johnston, J., Malnoy, M., Troggio, M., Perchepied, L., Sawyer, G., Wiedow, C., Won, K., Viola, R., Hellens, R.P., Brewer, L., Bus, V.G., Schaffer, R.J., Gardiner, S.E. and Velasco, R. (2014) The draft genome sequence of European pear (Pyrus communis L. ‘Bartlett’). PLoS One 9, e92644.

Chen, S., Zhou, Y., Chen, Y. and Gu, J. (2018) fastp: An ultra-fast all-in-one FASTQ preprocessor. Bioinformatics 34, i884–i890.

Cheng, H., Concepcion, G.T., Feng, X., Zhang, H. and Li, H. (2021) Haplotype-resolved de novo assembly using phased assembly graphs with hifiasm. Nat Methods 18, 170–175.

Conesa, A., Gotz, S., Garcia-Gomez, J.M., Terol, J., Talon, M. and Robles, M. (2005) Blast2GO: A universal tool for annotation, visualization and analysis in functional genomics research. Bioinformatics 21, 3674–3676.

Crow, T., Ta, J., Nojoomi, S., Aguilar-Rangel, M.R., Torres Rodriguez, J.V., Gates, D., Rellan-Alvarez, R., Sawers, R. and Runcie, D. (2020) Gene regulatory effects of a large chromosomal inversion in highland maize. PLoS Genet 16, e1009213.

Danecek, P., Auton, A., Abecasis, G., Albers, C.A., Banks, E., DePristo, M.A., Handsaker, R.E., Lunter, G., Marth, G.T. and Sherry, S.T. (2011) The variant call format and VCFtools. Bioinformatics 27, 2156–2158.

Ding, B., Liu, T., Hu, C., Song, Y., Hao, R., Feng, X., Cui, T., Han, Y. and Li, L. (2021) Comparative analysis of transcriptomic profiling to identify genes involved in the bulged surface of pear fruit (Pyrus bretschneideri Rehd. cv. Yuluxiangli). Physiol Mol Biol Plants 27, 69–80.

Dolatabadian, A., Yuan, Y., Bayer, P.E., Petereit, J., Severn-Ellis, A., Tirnaz, S., Patel, D., Edwards, D. and Batley, J. (2022) Copy number variation among resistance genes analogues in Brassica napus. Genes 13, 2037.

Dong, X., Wang, Z., Tian, L., Zhang, Y., Qi, D., Huo, H., Xu, J., Li, Z., Liao, R., Shi, M., Wahocho, S.A., Liu, C., Zhang, S., Tian, Z. and Cao, Y. (2020) De novo assembly of a wild pear (Pyrus betuleafolia) genome. Plant Biotechnol J 18, 581–595.

El Houari, I., Klima, P., Baekelandt, A., Staswick, P.E., Uzunova, V., Del Genio, C.I., Steenackers, W., Dobrev, P.I., Filepova, R., Novak, O., Napier, R., Petrasek, J., Inze, D., Boerjan, W. and Vanholme, B. (2023) Non-specific effects of the CINNAMATE-4-HYDROXYLASE inhibitor piperonylic acid. Plant J 115, 470–479.

Finn, R.D., Bateman, A., Clements, J., Coggill, P., Eberhardt, R.Y., Eddy, S.R., Heger, A., Hetherington, K., Holm, L., Mistry, J., Sonnhammer, E.L., Tate, J. and Punta, M. (2014) Pfam: the protein families database. Nucleic Acids Res 42, D222–230.

Flynn, J.M., Hubley, R., Goubert, C., Rosen, J., Clark, A.G., Feschotte, C. and Smit, A.F. (2020) RepeatModeler2 for automated genomic discovery of transposable element families. Proc Natl Acad Sci U S A 117, 9451–9457.

Gao, L., Gonda, I., Sun, H., Ma, Q., Bao, K., Tieman, D.M., Burzynski-Chang, E.A., Fish, T.L., Stromberg, K.A., Sacks, G.L., Thannhauser, T.W., Foolad, M.R., Diez, M.J., Blanca, J., Canizares, J., Xu, Y., van der Knaap, E., Huang, S., Klee, H.J., Giovannoni, J.J. and Fei, Z. (2019) The tomato pan-genome uncovers new genes and a rare allele regulating fruit flavor. Nat Genet 51, 1044–1051.

Gao, Y., Xu, J., Li, Z., Zhang, Y., Riera, N., Xiong, Z., Ouyang, Z., Liu, X., Lu, Z., Seymour, D., Zhong, B. and Wang, N. (2023) Citrus genomic resources unravel putative genetic determinants of Huanglongbing pathogenicity. iScience 26, 106024.

Goel, M., Sun, H., Jiao, W.B. and Schneeberger, K. (2019) SyRI: Finding genomic rearrangements and local sequence differences from whole-genome assemblies. Genome Biol 20, 277.

Golicz, A.A., Bayer, P.E., Barker, G.C., Edger, P.P., Kim, H., Martinez, P.A., Chan, C.K., Severn-Ellis, A., McCombie, W.R., Parkin, I.A., Paterson, A.H., Pires, J.C., Sharpe, A.G., Tang, H., Teakle, G.R., Town, C.D., Batley, J. and Edwards, D. (2016) The pangenome of an agronomically important crop plant Brassica oleracea. Nat Commun 7, 13390.

Guo, G., Wei, P., Yu, T., Zhang, H., Heng, W., Liu, L., Zhu, L. and Jia, B. (2022) PbrARF4 contributes to calyx shedding of fruitlets in ‘Dangshan Suli’pear by partly regulating the expression of abscission genes. Hortic Plant J 10.1016/j.hpj.2022.09.006

Haas, B.J., Salzberg, S.L., Zhu, W., Pertea, M., Allen, J.E., Orvis, J., White, O., Buell, C.R. and Wortman, J.R. (2008) Automated eukaryotic gene structure annotation using EVidenceModeler and the Program to Assemble Spliced Alignments. Genome Biol 9, R7.

Hamala, T., Wafula, E.K., Guiltinan, M.J., Ralph, P.E., dePamphilis, C.W. and Tiffin, P. (2021) Genomic structural variants constrain and facilitate adaptation in natural populations of Theobroma cacao, the chocolate tree. Proc Natl Acad Sci U S A 118, e2102914118

Holt, C. and Yandell, M. (2011) MAKER2: an annotation pipeline and genome-database management tool for second-generation genome projects. BMC Bioinformatics 12, 491.

Hu, H., Scheben, A., Verpaalen, B., Tirnaz, S., Bayer, P.E., Hodel, R.G.J., Batley, J., Soltis, D.E., Soltis, P.S. and Edwards, D. (2022) Amborella gene presence/absence variation is associated with abiotic stress responses that may contribute to environmental adaptation. New Phytol 233, 1548–1555.

Hu, H., Yuan, Y., Bayer, P.E., Fernandez, C.T., Scheben, A., Golicz, A.A. and Edwards, D. (2020) Legume pangenome construction using an iterative mapping and assembly approach. Methods Mol Biol 2107, 35–47.

Huerta-Cepas, J., Szklarczyk, D., Heller, D., Hernandez-Plaza, A., Forslund, S.K., Cook, H., Mende, D.R., Letunic, I., Rattei, T., Jensen, L.J., von Mering, C. and Bork, P. (2019) eggNOG 5.0: A hierarchical, functionally and phylogenetically annotated orthology resource based on 5090 organisms and 2502 viruses. Nucleic Acids Res 47, D309–D314.

Jeffares, D.C., Jolly, C., Hoti, M., Speed, D., Shaw, L., Rallis, C., Balloux, F., Dessimoz, C., Bähler, J. and Sedlazeck, F.J. (2017) Transient structural variations have strong effects on quantitative traits and reproductive isolation in fission yeast. Nat Commun 8, 14061.

Jiao, W.B. and Schneeberger, K. (2020) Chromosome-level assemblies of multiple Arabidopsis genomes reveal hotspots of rearrangements with altered evolutionary dynamics. Nat Commun 11, 989.

Jin, W., Wang, H., Li, M., Wang, J., Yang, Y., Zhang, X., Yan, G., Zhang, H., Liu, J. and Zhang, K. (2016) The R2R3 MYB transcription factor PavMYB10.1 involves in anthocyanin biosynthesis and determines fruit skin colour in sweet cherry (Prunus avium L.). Plant Biotechnol J 14, 2120–2133.

Kanehisa, M., Goto, S., Sato, Y., Furumichi, M. and Tanabe, M. (2012) KEGG for integration and interpretation of large-scale molecular data sets. Nucleic Acids Res 40, D109–114.

Korunes, K.L. and Samuk, K. (2021) pixy: Unbiased estimation of nucleotide diversity and divergence in the presence of missing data. Mol Ecol Resour 21, 1359–1368.

Langmead, B. and Salzberg, S.L. (2012) Fast gapped-read alignment with Bowtie 2. Nat Methods 9, 357–359.

Leger, A., Amaral, P.P., Pandolfini, L., Capitanchik, C., Capraro, F., Miano, V., Migliori, V., Toolan-Kerr, P., Sideri, T., Enright, A.J., Tzelepis, K., van Werven, F.J., Luscombe, N.M., Barbieri, I., Ule, J., Fitzgerald, T., Birney, E., Leonardi, T. and Kouzarides, T. (2021) RNA modifications detection by comparative Nanopore direct RNA sequencing. Nat Commun 12, 7198.

Letunic, I. and Bork, P. (2019) Interactive Tree Of Life (iTOL) v4: Recent updates and new developments. Nucleic Acids Res 47, W256–W259.

Li, H. (2013) Aligning sequence reads, clone sequences and assembly contigs with BWA-MEM. arXiv preprint arXiv:1303.3997.

Li, H. (2018) Minimap2: pairwise alignment for nucleotide sequences. Bioinformatics 34, 3094–3100.

Li, H., Wang, S., Chai, S., Yang, Z., Zhang, Q., Xin, H., Xu, Y., Lin, S., Chen, X., Yao, Z., Yang, Q., Fei, Z., Huang, S. and Zhang, Z. (2022) Graph-based pan-genome reveals structural and sequence variations related to agronomic traits and domestication in cucumber. Nat Commun 13, 682.

Li R, Bi R, Cai H, Zhao J, Sun P, Xu W, Zhou Y, Yang W, Zheng L, Chen XL, Wang G, Wang D, Liu J, Teng H, Li G. (2022) Melatonin functions as a broad-spectrum antifungal by targeting a conserved pathogen protein kinase. J Pineal Res 2022, 31, e12839.

Li, Y.H., Zhou, G., Ma, J., Jiang, W., Jin, L.G., Zhang, Z., Guo, Y., Zhang, J., Sui, Y., Zheng, L., Zhang, S.S., Zuo, Q., Shi, X.H., Li, Y.F., Zhang, W.K., Hu, Y., Kong, G., Hong, H.L., Tan, B., Song, J., Liu, Z.X., Wang, Y., Ruan, H., Yeung, C.K., Liu, J., Wang, H., Zhang, L.J., Guan, R.X., Wang, K.J., Li, W.B., Chen, S.Y., Chang, R.Z., Jiang, Z., Jackson, S.A., Li, R. and Qiu, L.J. (2014) De novo assembly of soybean wild relatives for pan-genome analysis of diversity and agronomic traits. Nat Biotechnol 32, 1045–1052.

Liu, Y., Du, H., Li, P., Shen, Y., Peng, H., Liu, S., Zhou, G.A., Zhang, H., Liu, Z., Shi, M., Huang, X., Li, Y., Zhang, M., Wang, Z., Zhu, B., Han, B., Liang, C. and Tian, Z. (2020) Pan-genome of wild and cultivated soybeans. Cell 182, 162–176 e113.

Love, M.I., Huber, W. and Anders, S. (2014) Moderated estimation of fold change and dispersion for RNA-seq data with DESeq2. Genome Biol 15, 550.

Lovell, J.T., Bentley, N.B., Bhattarai, G., Jenkins, J.W., Sreedasyam, A., Alarcon, Y., Bock, C., Boston, L.B., Carlson, J., Cervantes, K., Clermont, K., Duke, S., Krom, N., Kubenka, K., Mamidi, S., Mattison, C.P., Monteros, M.J., Pisani, C., Plott, C., Rajasekar, S., Rhein, H.S., Rohla, C., Song, M., Hilaire, R.S., Shu, S., Wells, L., Webber, J., Heerema, R.J., Klein, P.E., Conner, P., Wang, X., Grauke, L.J., Grimwood, J., Schmutz, J. and Randall, J.J. (2021) Four chromosome scale genomes and a pan-genome annotation to accelerate pecan tree breeding. Nat Commun 12, 4125.

Lowe, T.M. and Eddy, S.R. (1997) tRNAscan-SE: A program for improved detection of transfer RNA genes in genomic sequence. Nucleic Acids Res 25, 955–964.

Luo, M., Zhou, X., Sun, H., Zhou, Q., Ge, W., Sun, Y., Yao, M. and Ji, S. (2021) Insights into profiling of volatile ester and LOX-pathway related gene families accompanying post-harvest ripening of ‘Nanguo’pears. Food Chem 335, 127665.

Lyu, X., Xia, Y., Wang, C., Zhang, K., Deng, G., Shen, Q., Gao, W., Zhang, M, Liao, N., Ling, J., Bo, Y., Hu, Z., Yang, J., Zhang, M. (2023) Pan-genome analysis sheds light on structural variation-based dissection of agronomic traits in melon crops. Plant Physiol. 193, 1330–1348.

Ma, M., Liu, S., Wang, Z., Shao, R., Ye, J., Yan, W., Lv, H., Hasi, A. and Che, G. (2022) Genome-wide identification of the SUN gene family in melon (Cucumis melo) and functional characterization of two CmSUN genes in regulating fruit shape variation. Int J Mol Sci 23.

Marcais, G., Delcher, A.L., Phillippy, A.M., Coston, R., Salzberg, S.L. and Zimin, A. (2018) MUMmer4: A fast and versatile genome alignment system. PLoS Comput Biol 14, e1005944.

Mitchell, A., Chang, H.Y., Daugherty, L., Fraser, M., Hunter, S., Lopez, R., McAnulla, C., McMenamin, C., Nuka, G., Pesseat, S., Sangrador-Vegas, A., Scheremetjew, M., Rato, C., Yong, S.Y., Bateman, A., Punta, M., Attwood, T.K., Sigrist, C.J., Redaschi, N., Rivoire, C., Xenarios, I., Kahn, D., Guyot, D., Bork, P., Letunic, I., Gough, J., Oates, M., Haft, D., Huang, H., Natale, D.A., Wu, C.H., Orengo, C., Sillitoe, I., Mi, H., Thomas, P.D. and Finn, R.D. (2015) The InterPro protein families database: The classification resource after 15 years. Nucleic Acids Res 43, D213–221.

Nawrocki, E.P. and Eddy, S.R. (2013) Infernal 1.1: 100-fold faster RNA homology searches. Bioinformatics 29, 2933–2935.

O’Donovan, C., Martin, M.J., Gattiker, A., Gasteiger, E., Bairoch, A. and Apweiler, R. (2002) High-quality protein knowledge resource: SWISS-PROT and TrEMBL. Brief Bioinform 3, 275–284.

Ou, C., Wang, F., Wang, J., Li, S., Zhang, Y., Fang, M., Ma, L., Zhao, Y. and Jiang, S. (2019) A de novo genome assembly of the dwarfing pear rootstock Zhongai 1. Sci Data 6, 281.

Ou, S. and Jiang, N. (2018) LTR_retriever: A highly accurate and sensitive program for identification of long terminal repeat retrotransposons. Plant Physiol 176, 1410–1422.

Peat, T.S., Bottcher, C., Newman, J., Lucent, D., Cowieson, N. and Davies, C. (2012) Crystal structure of an indole-3-acetic acid amido synthetase from grapevine involved in auxin homeostasis. Plant Cell 24, 4525–4538.

Pencik, A., Simonovik, B., Petersson, S.V., Henykova, E., Simon, S., Greenham, K., Zhang, Y., Kowalczyk, M., Estelle, M., Zazimalova, E., Novak, O., Sandberg, G. and Ljung, K. (2013) Regulation of auxin homeostasis and gradients in Arabidopsis roots through the formation of the indole-3-acetic acid catabolite 2-oxindole-3-acetic acid. Plant Cell 25, 3858–3870.

Perina, A., Gonzalez-Tizon, A.M., Meilan, I.F. and Martinez-Lage, A. (2017) De novo transcriptome assembly of shrimp Palaemon serratus. Genom Data 11, 89–91.

Pertea, M., Kim, D., Pertea, G.M., Leek, J.T. and Salzberg, S.L. (2016) Transcript-level expression analysis of RNA-seq experiments with HISAT, StringTie and Ballgown. Nat Protoc 11, 1650–1667.

Pruitt, K.D., Tatusova, T. and Maglott, D.R. (2007) NCBI reference sequences (RefSeq): a curated non-redundant sequence database of genomes, transcripts and proteins. Nucleic Acids Res 35, D61–65.

Qi, X., Wu, J., Wang, L., Li, L., Cao, Y., Tian, L., Dong, X. and Zhang, S. (2013) Identifying the candidate genes involved in the calyx abscission process of ‘Kuerlexiangli’ (Pyrus sinkiangensis Yu) by digital transcript abundance measurements. BMC Genomics 14, 727.

Qian, M., Yu, B., Li, X., Sun, Y., Zhang, D. and Teng, Y. (2014) Isolation and expression analysis of anthocyanin biosynthesis genes from the red Chinese sand pear, Pyrus pyrifolia Nakai cv. Mantianhong, in response to methyl jasmonate treatment and UV-B/VIS conditions. Plant Mol Biol Rep 32, 428–437.

Qin, P., Lu, H., Du, H., Wang, H., Chen, W., Chen, Z., He, Q., Ou, S., Zhang, H., Li, X., Li, X., Li, Y., Liao, Y., Gao, Q., Tu, B., Yuan, H., Ma, B., Wang, Y., Qian, Y., Fan, S., Li, W., Wang, J., He, M., Yin, J., Li, T., Jiang, N., Chen, X., Liang, C. and Li, S. (2021) Pan-genome analysis of 33 genetically diverse rice accessions reveals hidden genomic variations. Cell 184, 3542–3558 e3516.

Raudvere, U., Kolberg, L., Kuzmin, I., Arak, T., Adler, P., Peterson, H. and Vilo, J. (2019) g:Profiler: A web server for functional enrichment analysis and conversions of gene lists (2019 update). Nucleic Acids Res 47, W191–W198.

Shi, F., Zhou, X., Yao, M.M., Zhou, Q., Ji, S.J. and Wang, Y. (2019) Low-temperature stress-induced aroma loss by regulating fatty acid metabolism pathway in ‘Nanguo’ pear. Food Chem 297, 124927.

Shirasawa, K., Itai, A. and Isobe, S. (2021) Chromosome-scale genome assembly of Japanese pear (Pyrus pyrifolia) variety ‘Nijisseiki’. DNA Res 28, dsab001.

Snouffer, A., Kraus, C. and van der Knaap, E. (2020) The shape of things to come: ovate family proteins regulate plant organ shape. Curr Opin Plant Biol 53, 98–105.

Song, J.M., Guan, Z., Hu, J., Guo, C., Yang, Z., Wang, S., Liu, D., Wang, B., Lu, S., Zhou, R., Xie, W.Z., Cheng, Y., Zhang, Y., Liu, K., Yang, Q.Y., Chen, L.L. and Guo, L. (2020) Eight high-quality genomes reveal pan-genome architecture and ecotype differentiation of Brassica napus. Nat Plants 6, 34–45.

Stanke, M., Keller, O., Gunduz, I., Hayes, A., Waack, S. and Morgenstern, B. (2006) AUGUSTUS: ab initio prediction of alternative transcripts. Nucleic Acids Res 34, W435–439.

Sun, X., Jiao, C., Schwaninger, H., Chao, C.T., Ma, Y., Duan, N., Khan, A., Ban, S., Xu, K. and Cheng, L. (2020) Phased diploid genome assemblies and pan-genomes provide insights into the genetic history of apple domestication. Nat genet 52, 1423–1432.

Tang, D., Jia, Y., Zhang, J., Li, H., Cheng, L., Wang, P., Bao, Z., Liu, Z., Feng, S., Zhu, X., Li, D., Zhu, G., Wang, H., Zhou, Y., Zhou, Y., Bryan, G.J., Buell, C.R., Zhang, C. and Huang, S. (2022) Genome evolution and diversity of wild and cultivated potatoes. Nature 606, 535–541.

Tarailo-Graovac, M. and Chen, N. (2009) Using RepeatMasker to identify repetitive elements in genomic sequences. Curr Protoc Bioinformatics **Chapter** 4, 4 10 11–14 10 14.

Tahir, U. Q., Zhu, X., Xing, F., Chen, L. (2019) ppsPCP: A plant presence/absence variants scanner and pan-genome construction pipeline. Bioinformatics 35, 4156–4158.

Tahir, U. Q., Zhu, X., Khan, M.S., Xing, F., Chen, L. (2020) Pan-genome: A promising resource for noncoding RNA discovery in plants. Plant Genome 13, e20046.

Tie W, Zhou, F., Wang, L., Xie, W., Chen, H., Li, X., Lin, Y. (2012) Reasons for lower transformation efficiency in indica rice using Agrobacterium tumefaciens-mediated transformation: lessons from transformation assays and genome-wide expression profiling. Plant Mol Biol. 78, 1–18.

Torkamaneh, D., Lemay, M.A. and Belzile, F. (2021) The pan-genome of the cultivated soybean (PanSoy) reveals an extraordinarily conserved gene content. Plant Biotechnol J 19, 1852–1862.

Varshney, R.K., Roorkiwal, M., Sun, S., Bajaj, P., Chitikineni, A., Thudi, M., Singh, N.P., Du, X., Upadhyaya, H.D., Khan, A.W., Wang, Y., Garg, V., Fan, G., Cowling, W.A., Crossa, J., Gentzbittel, L., Voss-Fels, K.P., Valluri, V.K., Sinha, P., Singh, V.K., Ben, C., Rathore, A., Punna, R., Singh, M.K., Tar’an, B., Bharadwaj, C., Yasin, M., Pithia, M.S., Singh, S., Soren, K.R., Kudapa, H., Jarquin, D., Cubry, P., Hickey, L.T., Dixit, G.P., Thuillet, A.C., Hamwieh, A., Kumar, S., Deokar, A.A., Chaturvedi, S.K., Francis, A., Howard, R., Chattopadhyay, D., Edwards, D., Lyons, E., Vigouroux, Y., Hayes, B.J., von Wettberg, E., Datta, S.K., Yang, H., Nguyen, H.T., Wang, J., Siddique, K.H.M., Mohapatra, T., Bennetzen, J.L., Xu, X. and Liu, X. (2021) A chickpea genetic variation map based on the sequencing of 3,366 genomes. Nature 599, 622–627.

Walkowiak, S., Gao, L., Monat, C., Haberer, G., Kassa, M.T., Brinton, J., Ramirez-Gonzalez, R.H., Kolodziej, M.C., Delorean, E., Thambugala, D., Klymiuk, V., Byrns, B., Gundlach, H., Bandi, V., Siri, J.N., Nilsen, K., Aquino, C., Himmelbach, A., Copetti, D., Ban, T., Venturini, L., Bevan, M., Clavijo, B., Koo, D.H., Ens, J., Wiebe, K., N’Diaye, A., Fritz, A.K., Gutwin, C., Fiebig, A., Fosker, C., Fu, B.X., Accinelli, G.G., Gardner, K.A., Fradgley, N., Gutierrez-Gonzalez, J., Halstead-Nussloch, G., Hatakeyama, M., Koh, C.S., Deek, J., Costamagna, A.C., Fobert, P., Heavens, D., Kanamori, H., Kawaura, K., Kobayashi, F., Krasileva, K., Kuo, T., McKenzie, N., Murata, K., Nabeka, Y., Paape, T., Padmarasu, S., Percival-Alwyn, L., Kagale, S., Scholz, U., Sese, J., Juliana, P., Singh, R., Shimizu-Inatsugi, R., Swarbreck, D., Cockram, J., Budak, H., Tameshige, T., Tanaka, T., Tsuji, H., Wright, J., Wu, J., Steuernagel, B., Small, I., Cloutier, S., Keeble-Gagnere, G., Muehlbauer, G., Tibbets, J., Nasuda, S., Melonek, J., Hucl, P.J., Sharpe, A.G., Clark, M., Legg, E., Bharti, A., Langridge, P., Hall, A., Uauy, C., Mascher, M., Krattinger, S.G., Handa, H., Shimizu, K.K., Distelfeld, A., Chalmers, K., Keller, B., Mayer, K.F.X., Poland, J., Stein, N., McCartney, C.A., Spannagl, M., Wicker, T. and Pozniak, C.J. (2020) Multiple wheat genomes reveal global variation in modern breeding. Nature 588, 277–283.

Wang, P., Wu, X., Shi, Z., Tao, S., Liu, Z., Qi, K., Xie, Z., Qiao, X., Gu, C., Yin, H., Cheng, M., Gu, X., Liu, X., Tang, C., Cao, P., Xu, S., Zhou, B., Gu, T., Bian, Y., Wu, J. and Zhang, S. (2023a) A large-scale proteogenomic atlas of pear. Mol Plant 16, 599–615.

Wang, J., Yang, W., Zhang, S., Hu, H., Yuan, Y., Dong, J., Chen, L., Ma, Y., Yang, T., Zhou, L., Chen, J., Liu, B., Li, C., Edwards, D. and Zhao, J. (2023b) A pangenome analysis pipeline provides insights into functional gene identification in rice. Genome Biol 24, 19.

Wang, J., Hu, H., Liang, X., Tahir, Ul Qamar M., Zhang, Y., Zhao, J., Ren, H., Yan, X., Ding, B., Guo, J. (2023c) High-quality genome assembly and comparative genomic profiling of yellowhorn (Xanthoceras sorbifolia) revealed environmental adaptation footprints and seed oil contents variations. Front Plant Sci 14, 1147946.

Wang, X., Du, J. and Yao, X. (2015) Structural and dynamic basis of acid amido synthetase GH3.1: an investigation of substrate selectivity and major active site access channels. Mol Biosyst 11, 809–818.

Wang, Y., Strauss, S., Liu, S., Pieper, B., Lymbouridou, R., Runions, A. and Tsiantis, M. (2022) The cellular basis for synergy between RCO and KNOX1 homeobox genes in leaf shape diversity. Curr Biol 32, 3773–3784. e3775.

Wright, M.N., Gola, D. and Ziegler, A. (2017) Preprocessing and quality control for whole-genome sequences from the illumina HiSeq X platform. Methods Mol Biol 1666, 629–647.

Wu, B., Gao, L., Gao, J., Xu, Y., Liu, H., Cao, X., Zhang, B. and Chen, K. (2017) Genome-wide identification, expression patterns, and functional analysis of UDP glycosyltransferase family in peach (Prunus persica L. Batsch). Front Plant Sci 8, 389.

Wu, J., Wang, Y., Xu, J., Korban, S.S., Fei, Z., Tao, S., Ming, R., Tai, S., Khan, A.M., Postman, J.D., Gu, C., Yin, H., Zheng, D., Qi, K., Li, Y., Wang, R., Deng, C.H., Kumar, S., Chagne, D., Li, X., Wu, J., Huang, X., Zhang, H., Xie, Z., Li, X., Zhang, M., Li, Y., Yue, Z., Fang, X., Li, J., Li, L., Jin, C., Qin, M., Zhang, J., Wu, X., Ke, Y., Wang, J., Yang, H. and Zhang, S. (2018) Diversification and independent domestication of Asian and European pears. Genome Biol 19, 77.

Wu, J., Wang, Z., Shi, Z., Zhang, S., Ming, R., Zhu, S., Khan, M.A., Tao, S., Korban, S.S., Wang, H., Chen, N.J., Nishio, T., Xu, X., Cong, L., Qi, K., Huang, X., Wang, Y., Zhao, X., Wu, J., Deng, C., Gou, C., Zhou, W., Yin, H., Qin, G., Sha, Y., Tao, Y., Chen, H., Yang, Y., Song, Y., Zhan, D., Wang, J., Li, L., Dai, M., Gu, C., Wang, Y., Shi, D., Wang, X., Zhang, H., Zeng, L., Zheng, D., Wang, C., Chen, M., Wang, G., Xie, L., Sovero, V., Sha, S., Huang, W., Zhang, S., Zhang, M., Sun, J., Xu, L., Li, Y., Liu, X., Li, Q., Shen, J., Wang, J., Paull, R.E., Bennetzen, J.L., Wang, J. and Zhang, S. (2013) The genome of the pear (Pyrus bretschneideri Rehd.). Genome Res 23, 396–408.

Wu, S., Sun, H., Gao, L., Branham, S., McGregor, C., Renner, S.S., Xu, Y., Kousik, C., Wechter, W.P., Levi, A. and Fei, Z. (2023) A Citrullus genus super-pangenome reveals extensive variations in wild and cultivated watermelons and sheds light on watermelon evolution and domestication. Plant Biotechnol J 21, 1926–1928.

Wu, X., Shi, X., Bai, M., Chen, Y., Li, X., Qi, K., Cao, P., Li, M., Yin, H. and Zhang, S. (2019) Transcriptomic and gas chromatography-mass spectrometry metabolomic profiling analysis of the epidermis provides insights into cuticular wax regulation in developing ‘Yuluxiang’ Pear Fruit. J Agric Food Chem 67, 8319–8331.

Yang, J., Lee, S.H., Goddard, M.E. and Visscher, P.M. (2011) GCTA: A tool for genome-wide complex trait analysis. Am J Hum Genet 88, 76–82.

Yang, T., Liu, R., Luo, Y., Hu, S., Wang, D., Wang, C., Pandey, M.K., Ge, S., Xu, Q. and Li, N. (2022) Improved pea reference genome and pan-genome highlight genomic features and evolutionary characteristics. Nat genet 54, 1553–1563.

Yu, J., Golicz, A.A., Lu, K., Dossa, K., Zhang, Y., Chen, J., Wang, L., You, J., Fan, D. and Edwards, D. (2019) Insight into the evolution and functional characteristics of the pan-genome assembly from sesame landraces and modern cultivars. Plant Biotechnol J 17, 881–892.

Zhang, B., Li, Q., Keyhaninejad, N., Taitano, N., Sapkota, M., Snouffer, A. and van der Knaap, E. (2023) A combinatorial TRM-OFP module bilaterally fine-tunes tomato fruit shape. New Phytol 238, 2393–2409.

Zhang, B., Shen, J.Y., Wei, W.W., Xi, W.P., Xu, C.J., Ferguson, I. and Chen, K. (2010) Expression of genes associated with aroma formation derived from the fatty acid pathway during peach fruit ripening. J Agric Food Chem 58, 6157–6165.

Zhang, J., Jiang, H., Li, Y., Wang, S., Wang, B., Xiao, J. and Cao, Y. (2022) Transcriptomic and physiological analysis reveals the possible mechanism of ultrasound inhibiting strawberry (Fragaria x ananassa Duch.) postharvest softening. Front Nutr 9, 1066043.

Zhang, M.Y., Xue, C., Xu, L., Sun, H., Qin, M.F., Zhang, S. and Wu, J. (2016) Distinct transcriptome profiles reveal gene expression patterns during fruit development and maturation in five main cultivated species of pear (Pyrus L.). Sci Rep 6, 28130.

Zhao, J., Bayer, P.E., Ruperao, P., Saxena, R.K., Khan, A.W., Golicz, A.A., Nguyen, H.T., Batley, J., Edwards, D. and Varshney, R.K. (2020) Trait associations in the pangenome of pigeon pea (Cajanus cajan). Plant Biotechnol J 18, 1946–1954.

Zhou, Y., Zhang, Z., Bao, Z., Li, H., Lyu, Y., Zan, Y., Wu, Y., Cheng, L., Fang, Y., Wu, K., Zhang, J., Lyu, H., Lin, T., Gao, Q., Saha, S., Mueller, L., Fei, Z., Stadler, T., Xu, S., Zhang, Z., Speed, D. and Huang, S. (2022) Graph pangenome captures missing heritability and empowers tomato breeding. Nature 606, 527–534.

Zhou, Y., Minio, A., Massonnet, M., Solares, E., Lv, Y., Beridze, T., Cantu, D., Gaut, BS. (2019) The population genetics of structural variants in grapevine domestication. Nat Plants 5, 965–979.

Zmienko, A., Samelak, A., Kozlowski, P. and Figlerowicz, M. (2014) Copy number polymorphism in plant genomes. Theor Appl Genet 127, 1–18.

